# Mapping the functional impact of non-coding regulatory elements in primary T cells through single-cell CRISPR screens

**DOI:** 10.1101/2023.05.14.540711

**Authors:** Celia Alda Catalinas, Ximena Ibarra-Soria, Christina Flouri, Jorge Esparza Gordillo, Diana Cousminer, Anna Hutchinson, Adam Krejci, Adrian Cortes, Alison Acevedo, Sunir Malla, Carl Fishwick, Gerard Drewes, Radu Rapiteanu

## Abstract

Drug targets with human genetic evidence are expected to increase clinical success by at least two-fold. Yet, translating disease-associated genetic variants into functional knowledge remains a fundamental challenge of early drug discovery. A key issue is that, currently, the vast majority of complex disease associations cannot be cleanly mapped to a gene. Immune disease-associated variants are enriched within regulatory elements, such as distal enhancers, found in T cell-specific open chromatin regions. To identify the genes and thus the molecular programs modulated by these regulatory elements, we developed a CRISPRi-based single-cell functional screening approach in primary human CD4^+^ T cells. Our pipeline enables the interrogation of transcriptomic changes induced by the perturbation of regulatory elements at scale. We first optimised a highly efficient CRISPRi protocol in primary human CD4^+^ T cells via CROPseq vectors. Subsequently, we performed a proof-of-concept screen targeting 45 non-coding regulatory elements and 35 transcription start sites and profiled approximately 250,000 CD4^+^ T cell single-cell transcriptomes. We developed a bespoke analytical pipeline for element-to-gene (E2G) mapping and demonstrate that our method can identify both previously annotated and novel E2G links. Lastly, we integrated genetic association data for immune-related traits and demonstrate how our platform can aid in the identification of effector genes for GWAS loci.

## Introduction

Genome-wide association studies (GWAS) have revealed thousands of disease-associated single-nucleotide polymorphisms (SNPs) (Buniello et al. 2019). Drug targets supported by human genetic evidence are expected to increase clinical success by at least two-fold (Nelson et al. 2015, Finan et al. 2017, Pritchard et al. 2017, King et al. 2019, Ochoa et al. 2022). Thus, understanding the molecular mechanisms underpinning GWAS hits is key to reducing attrition in drug discovery. More than 90% of disease-associated variants are located in non-coding genomic regions (Maurano et al. 2012, GTEX-Consortium 2020, Alsheikh et al. 2022), making it challenging to identify the causal effector gene(s) they regulate (Giambartolomei et al. 2014, Montefiori et al. 2018, French and Edwards 2020, Nasser et al. 2021, Umans et al. 2021).

Non-coding disease-associated variants are enriched within cell type-specific open chromatin regions, especially regulatory elements such as promoters and enhancers (Degner et al. 2012, ENCODE-Project-Consortium 2012, Maurano et al. 2012, Kundaje et al. 2015, Finucane et al. 2018, Montefiori et al. 2018, Calderon et al. 2019, Nott et al. 2019, Meuleman et al. 2020), and they often impact gene expression in a cell type-specific manner (Mumbach et al. 2017, Zhernakova et al. 2017, Schmiedel et al. 2018, Donovan et al. 2020, Kim-Hellmuth et al. 2020, Chiou et al. 2021, Ota et al. 2021, Young et al. 2021, Yazar et al. 2022). Hence, several studies have combined genetic fine-mapping with epigenomic profiles to prioritise candidate *cis*-regulatory elements within trait-relevant cell populations (Trynka and Raychaudhuri 2013, Farh et al. 2015, Gaulton et al. 2015, Chen et al. 2016, Soskic et al. 2019, Ulirsch et al. 2019, Amariuta et al. 2020, Moore et al. 2020, Boix et al. 2021). However, identifying the genes and downstream molecular programs modulated by disease-associated regulatory elements remains difficult with currently available tools.

CD4^+^ T cells play critical roles in autoimmune and inflammatory disorders, such as inflammatory bowel disease, type 1 diabetes, Crohn’s disease and rheumatoid arthritis (Skapenko et al. 2005, Sakaguchi et al. 2020). These cells are heterogeneous and highly plastic as they differentiate, and acquire distinct functions to counter pathogens and navigate changing environments (Zhu et al. 2010). Fine-mapping of GWAS loci, expression quantitative trait loci (eQTLs) and epigenomic studies have shown that immune disease-associated risk variants are enriched in CD4^+^ T cell regulatory regions (Trynka and Raychaudhuri 2013, Farh et al. 2015, Mumbach et al. 2017, Finucane et al. 2018, Calderon et al. 2019, Soskic et al. 2019, Nathan et al. 2021, Ota et al. 2021, Bossini-Castillo et al. 2022).

CRISPR is a powerful tool to functionally characterize and map non-coding regulatory elements to genes (Klann et al. 2017, Simeonov et al. 2017, Xie et al. 2017, Fulco et al. 2019, Gasperini et al. 2019, Gasperini et al. 2020, Freimer et al. 2021, Morris et al. 2023). In recent years, the combination of CRISPR screening with single-cell RNA-sequencing (scRNA-seq) has enabled deep phenotyping of genetic perturbations at scale (Shifrut et al. 2018, Gate et al. 2019, Schumann et al. 2020), providing an unprecedented opportunity to disentangle genome regulation. Specifically, high-throughput single-cell CRISPR-interference (CRISPRi) screens of regulatory regions have been performed to generate enhancer-gene maps at scale (Gasperini et al. 2019, Morris et al. 2023). However, these studies have so far used immortalised cell lines due to their ease of manipulation. However, the epigenetic regulation of immortalised, highly passaged cell lines is adapted to their highly proliferative state rather than being representative of the tissues from which they were derived (Pastor et al. 2010, Kaur and Dufour 2012). Therefore, we sought to establish a method that allowed us to query the function of regulatory elements in a physiologically relevant cell context.

Here, we present the method called “primary T cell crisprQTL”, a high-throughput, single-cell, pooled CRISPRi-based functional screening framework to map non-coding regulatory elements to genes in primary human CD4^+^ T cells. We perturbed transcription start sites (TSSs) and putative regulatory elements with ZIM3-dCas9 and profiled 250,195 high-quality single CD4^+^ T cell transcriptomes. We developed an analytical pipeline to robustly assign perturbations to cells and to determine gene expression changes. We demonstrate that our method can identify high-confidence *cis* element-to-gene (E2G) pairs and nominate novel E2G links supported by genetic evidence.

## Results

### Implementation of single-cell CRISPRi technology in human primary CD4^+^ T cells

To enable lentiviral-based CRISPRi perturbations in unexpanded primary CD4^+^ T cells, we first targeted the TSS of the *CD4* gene using an SFFV-dCas9-KOX1 construct. Briefly, T cells were activated and transduced with dCas9-KOX1 lentivirus and selected with blasticidin. Following selection, cells were re-activated and transduced with a *CD4* TSS-targeting guide RNA (gRNA) or with a non-targeting (NT) control, cloned into a CROPseq (CRISPR droplet sequencing) (Datlinger et al. 2017) backbone (see Materials and Methods). After puromycin selection of cells expressing the gRNA, the effect of the CRISPRi perturbation on the expression of CD4 protein was analysed by flow cytometry (Fig. 1A,B). This initial experiment resulted in 75% downregulation of CD4 at day 10 post-gRNA transduction (Fig. 1B).

**Figure 1:**
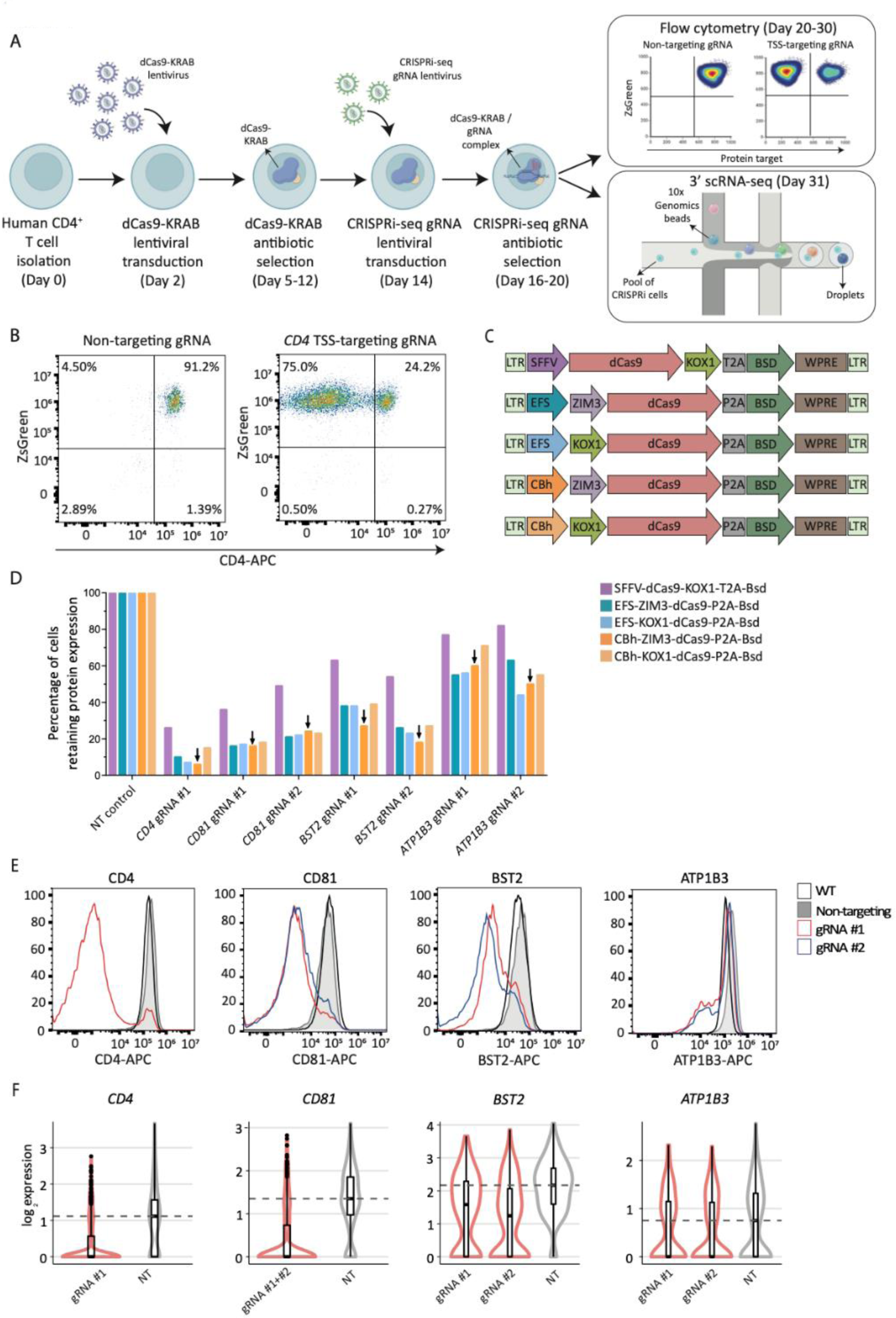
**A)** Schematic of the CRISPRi protocol in primary CD4^+^ T cells. **B)** Flow cytometry scatterplot showing CD4 expression (CD4-APC staining, *x-axis*) vs ZsGreen expression(reporter in gRNA lentiviral construct, *y-axis*) in primary CD4^+^ T cells expressing SFFV-dCas9-KOX1, transduced with a non-targeting gRNA control or a CD4 TSS-targeting gRNA. Cells were selectedwith puromycinand analysed 10 days after gRNA transduction. **C)** Schematic of dCas9-repressor lentiviral constructs tested. **D)** Bar plot showing the percentage of cells retaining protein expression for different target genes (*CD4, CD81, BST2, ATP1B3*) 10 days after TSS-targeting gRNA transduction into primary CD4^+^ T cells expressing different dCas9-repressor constructs, analysed by flow cytometry and normalised to the corresponding non-targeting gRNA control sample. gRNA #1 and #2 refer to two different gRNA designs for a given TSS. **E)** Histograms showing expression of the target gene (*CD4, CD81, BST2, ATP1B3*) 10 days after gRNA transduction into primary CD4^+^ T cells expressing a CBh-ZIM3-dCas9 repressor construct, analysed by flow cytometry. The wild-type (WT) control are non-transduced cells stained with the same antibody for the corresponding target gene. **F)** Normalised expression levels of the same target genes, measured by 10X Genomics 3’ scRNA-seq 11 days after the corresponding targeting (red) or non-targeting (grey) gRNAs were transduced into primary CD4^+^ T cells expressing a CBh-ZIM3-dCas9 repressor construct. Dashed line indicates median expression level in cells with non-targeting controls. Note gRNA #1 and #2 for *CD81* TSS were analysed together due to sequence similarity.

To optimise CRISPRi efficiency and assess gene-to-gene targeting variability, we tested a panel of dCas9-repressor lentiviral constructs targeting the TSS of four cell surface receptors: *CD4*, *CD81*, *BST2* and *ATP1B3*. We designed dCas9 constructs using three repressor domains (KOX1, ZIM3, and the fusion KOX1-MeCP2) under the control of three promoters (SFFV, EFS, and CBh) (Fig. 1C). The KRAB domain ZIM3 and the fusion of KOX1-MeCP2 have been shown to improve silencing efficiency compared to the more widely used KOX1 domain (Alerasool et al. 2020). Following the workflow described above (Fig. 1A), the effect of CRISPRi perturbations on the expression of the four cell surface receptors was assessed by flow cytometry at days 6, 10 and 16 post-transduction. We achieved successful transduction and blasticidin selection for all repressor constructs except for SFFV-dCas9-KOX1-MeCP2, likely due to the large provirus size (see Materials and Methods). We observed variable silencing efficacy for different genes (Fig. 1D-E), likely due to differences in gRNA efficiency, basal gene expression and local chromatin context, consistent with previous CRISPR modulation studies (Gilbert et al. 2014, Alda-Catalinas et al. 2020). Importantly, expression of the dCas9-repressor domain under the CBh or EFS promoters led to enhanced downregulation of the targeted genes, compared to the SFFV promoter (e.g., 84.7%-93.1% CD4 downregulation with CBhor EFS vs 73.4% CD4 downregulation with SFFV-dCas9-KOX1) (Fig. 1D). Generally, using a ZIM3 repressor under a CBh promoter resulted in improved silencing compared to a KOX1 repressor across multiple target genes (e.g., 93.1% vs 84.7% downregulation for CD4, 77% vs 66.7% for BST2 and 44.3% vs 36.2% for ATP1B3; average of two gRNAs per target) (Fig. 1D-E) and timepoints (Fig. S1A). Thus, we selected the CBh-ZIM3-dCas9 construct for all subsequent experiments.

Next, we assessed whether we could detect gRNA transcripts and quantify the downregulation of targetedgenes by single-cell transcriptomics. Following independent transductions of gRNAs targeting the TSS of *CD4*, *CD81*, *BST2* and *ATP1B3*, we pooled all cells and performed a CROPseq experiment using 10X Genomics 3’ chemistry. We were able to confidently assign gRNAs to cells (see Materials and Methods) and observed significant target gene downregulation compared to non-targeting controls (Fig. 1F). Additionally, the magnitude of the expression changes measured by scRNA-seq were highly correlated with the protein abundance changes detected by flow cytometry (Fig. S1B). These results show high-efficiency CRISPRi-based silencing of gene expression in primary CD4^+^ T cells and demonstrate that perturbation effects can be read-out at single-cell resolution using a CROPseq-based approach.

### Proof-of-concept crisprQTL screen recapitulates previously validated element-to-gene links

Having established the CRISPRi CROPseq workflow in primary T cells, we sought to use this pipeline for E2G mapping. We refer to this method as primary T cell crisprQTL. To show proof-of-concept, we silenceda panel of non-coding elements likely to regulate gene expression in primary T cells, alongside technical controls (Fig. 2A). First, we selected the *CD2* locus control region (LCR), which contains three regulatory elements that enhance *CD2* gene expression (Lake et al. 1990, Kaptein et al. 1998). Second, we identified 28 enhancers that overlap open chromatin in primary CD4^+^ T cells and were previously paired to a gene in a functional E2G mapping study in the K562 cell line (leukaemia bone marrow-isolated lymphoblast cells) (Gasperini et al. 2019) (see Materials and Methods); we refer to these elements as Gasperini enhancers (Gasperini_ENH). Third, we selected 14 non-coding elements from the ENCODE registry of candidate *cis*-Regulatory Elements (cCREs), five of which were intergenic while the other nine overlapped introns. Lastly, we included 35 TSSs of genes spanning a wide range of gene expression levels to serve as technical controls (Fig. 2A). For most of these perturbations, there is a predicted or *expected* downregulated gene based on previous evidence: *CD2* for LCR perturbations; the gene linked in K562 cells for enhancers selected from Gasperini et al. (Gasperini et al. 2019); and the gene immediately downstream of the TSS. Thus, we treated these perturbations as positive controls, whereas the cCRE perturbations were selected for exploratory E2G mapping.

**Figure 2:**
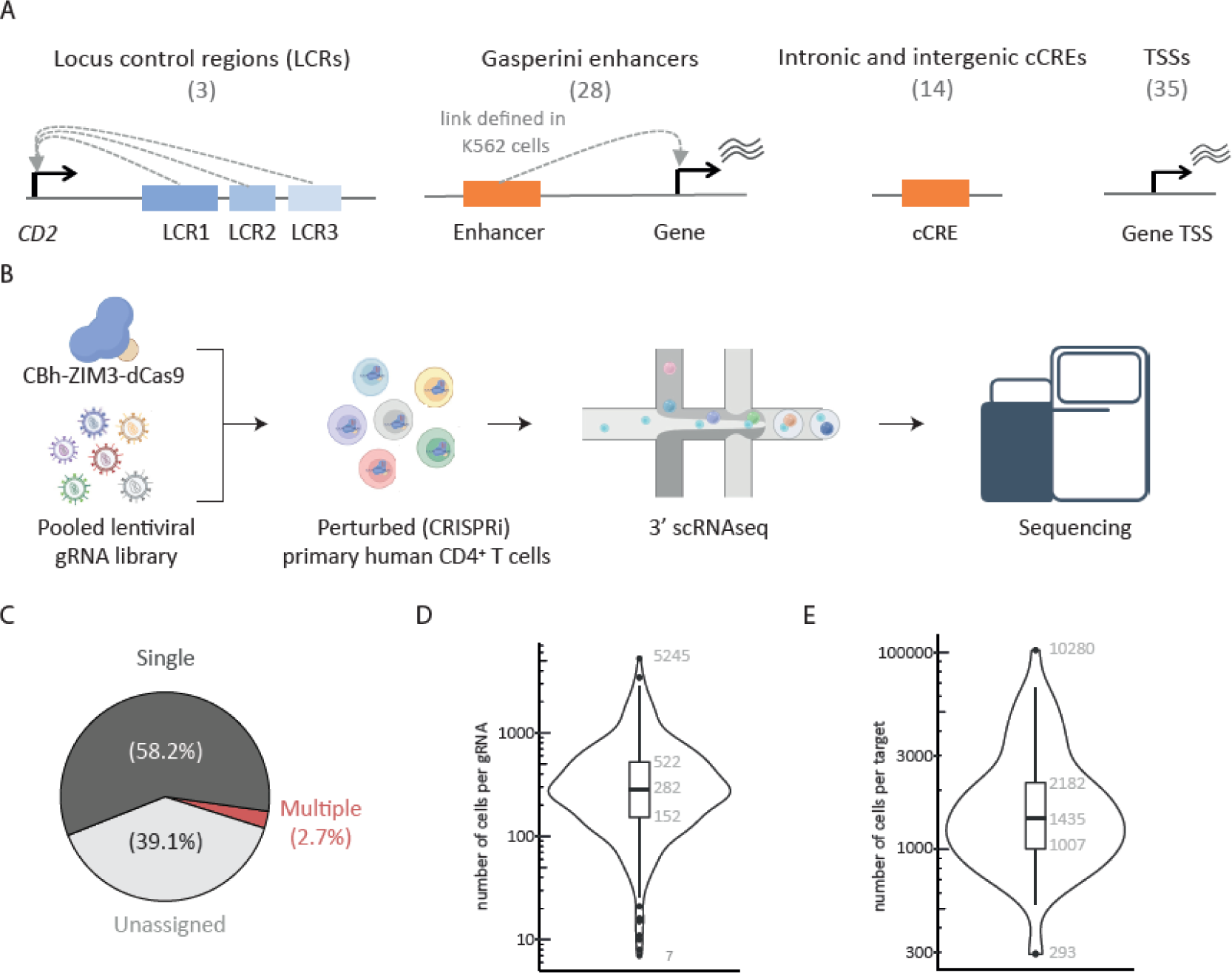
**A)** Schematic of the classes of loci targeted in the crisprQTL screen, including the locus control regions of *CD2*, enhancers linked to genes from Gasperini *et al*. (2019), regulatory elements (intronic and intergenic) overlapping ENCODE cCREs, and gene transcription start sites (TSS). **B)** Schematic of primary T cell crisprQTL experimental approach: CBh-ZIM3-dCas9 and the pooled gRNA library were introduced as described in Fig. 1A, and perturbed cells were analysed by 10X Genomics 3’ scRNA-seq. **C)** Proportion of cells from the screen with a single gRNA, multiple gRNAs, or none (unassigned). **D)** Distribution of the number of cells recovered with each gRNA in the pooled library. Numbers indicate, from top to bottom, the maximum, 75 ^th^, 50^th^, 25^th^ quantiles, and minimum. **E)** Same as D but for the number of cells per target (each target is targeted by four gRNAs).

We designed four gRNAs to target each candidate regulatory element or TSS along with 35 non-targeting controls (Table S1) and cloned the resulting 355 gRNA library into our CRISPRi CROPseq backbone. This pooled gRNA library was transduced at low multiplicity of infection (MOI) into primary CD4^+^ T cells previously selected for CBh-ZIM3-dCas9 expression. We generated high-quality transcriptional profiles and gRNA amplicon libraries for 250,195 single cells using the 3’ scRNA-seq 10X Genomics platform (Fig. 2B, S2A).

To assign gRNAs to cells, we developed a probabilistic framework based on a binomial distribution to assess the likelihood that a gRNA is present in a cell, taking into consideration its representation in the initial plasmid library. Unlike the static thresholding methods previously employed (Hill et al. 2018, Gasperini et al. 2019), we found that our framework allows us to control for two important sources of technical noise: biases in the efficiency of gRNA transcript recovery between cells, and variation in the abundance of each gRNA species in the experiment, which influences the quantification noise from ambient RNA and PCR artefacts. After applying this method to our gRNA amplicon data, we confidently assigned at least one gRNA to 152,403 (61%) cells (Fig. 2C, S2B). Importantly, we verified that these gRNA assignments were consistent with gRNA transcripts recovered in the gene expression library, confirming that the PCR enrichment process does not introduce spurious signals (Fig. S2C). As cells were transduced at low MOI, we identified a single gRNA in most cells (95.6% of cells assigned) (Fig. 2C). We recovered all 355 gRNAs present in the library, with a median of 282 cells per gRNA (Fig. 2D). All but 19 gRNAs were identified in more than 50 cells, and the numbers of cells assigned to each perturbation were correlated with the gRNA abundances in the plasmid library, indicating no significant adverse effects on cell fitness or viability (Fig. S2D). The gRNAs showing poor recovery corresponded to a variety of targets. Thus, we achieved good representation of all perturbations in the experiment (median of 1,435 cells per target; Fig. 2E).

To determine the effects of each perturbation on the expression of nearby genes (1 Mb up and downstream from the targeting site) we used MAST, an algorithm for scRNA-seq differential expression analysis (Finak et al. 2015). We first looked at the *expected* gene in the positive control perturbations and excluded two TSS and two Gasperini enhancer positive control targets with poor expression (detection in fewer than 5% of cells). Cells transduced with gRNAs targeting a TSS showed significant (FDR < 5%) downregulation for 32 out of the 33 targets (Fig. 3A-C), with an average reduction of 31% from wild-type expression levels (effect sizes in the range of 3%-97%; Fig. 3B). Similarly, *CD2* expression was significantly downregulated upon silencing of any of the three LCR regions targeted (average 29% reduction; Fig. 3A-C). For the E2G pairs identified by Gasperini et al. in K562 cells (Gasperini et al. 2019) (Gasperini_ENH), we reproduced 17 out of the 26 associations in our primary T cell data, with an average reduction in gene expression levels of 20% (range 5%-50%; Fig. 3A-C).

**Figure 3:**
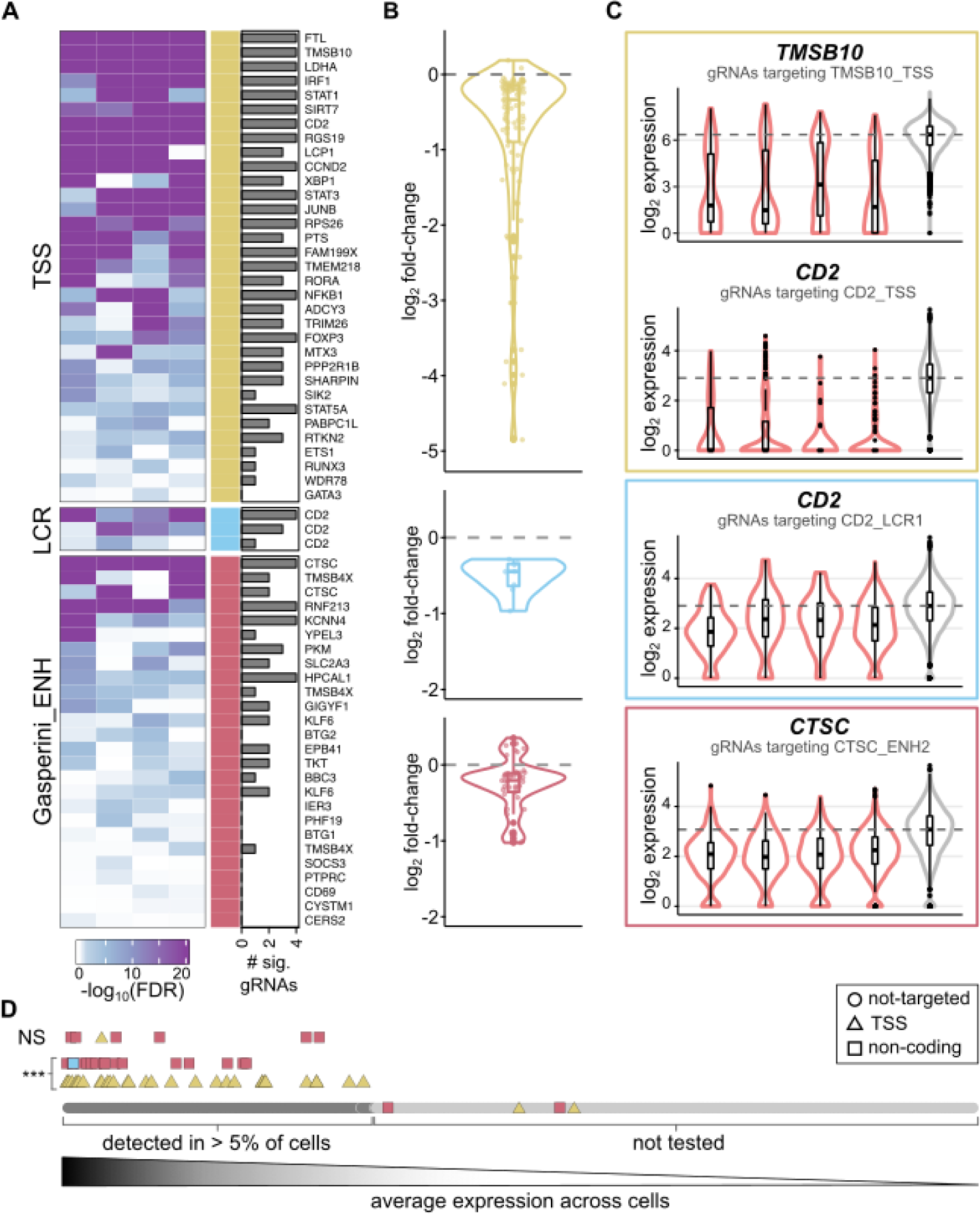
**A)** Heatmap of the differential expression significance values (−log_10_ adjusted p-value) for each of the four gRNAs targeting eachof the positive controlperturbations, whencomparing the expression of the expected gene in perturbed cells versus non-targeting controls. Different classes of targets are indicated by coloured bars (TSS - yellow, LCR - blue and Gasperini enhancers - red). The barplots to the right indicatehow many of the four gRNAs reach statistical significance. **B)** Distributions of the log_2_ fold-change values for all expected genes from positive control targets, split by target class. **C)** Representative examples of targets from each class. Normalised expression values in cells with targeting gRNAs (red) versus NT controls (grey) are shown. The title of the plot indicates the gene plotted. **D)** Plot depicting the effect of gene expression levels on our ability to detect downregulation effects upon perturbationof TSS and non-coding targets. At the bottom, all genes in the human genome are ranked by decreasing average expression in the scRNA-seq dataset. Only genes detected in at least 5% of the cells (dark grey) were considered in the differential expressionanalyses. Non testedgenes (light grey) include both genes not expressed in T cells, and genes not detected by scRNA-seq. Then, *expected* genes in positive control perturbations that were significantlydifferentially expressed(***) are indicated, separately for TSS (yellow triangles) and non-coding control perturbations (red squares for Gasperini_ENH target genes, blue square for *CD2*). Above, *expected* genes that were detected but not recovered as significantly downregulated upon perturbation(NS).

For gRNAs detected in at least 30 cells, we observed two major limitations in our ability to detect significant gene expression changes: the magnitude of the effect and the expression level of the affected gene (Fig. 3B, 3D). Since silencing the promoter results in strong downregulation of gene expression, we were able to detect the effects from TSS perturbations for genes covering the whole dynamic range of expression levels detected by single-cell transcriptomics (Fig. 3D). In contrast, silencing non-coding elements that affect genes expressed at lower levels, did not reach statistical significance (Fig. 3D). Importantly, targets with significant effects were supported by at least two different gRNAs in 83% of cases, for both promoter and non-coding elements (Fig. 3A). Altogether, these results demonstrate that our crisprQTL method efficiently silences both gene promoters and distal regulatory elements, and that the effects of these perturbations on gene expression can be detected by single-cell transcriptomics, enabling the mapping of regulatory element to genes in primary human CD4^+^ T cells.

### Evaluation of analytical methods for high confidence *cis* E2G mapping using crisprQTL

Having shown proof-of-concept based on downregulation of the *expected* gene for positive control perturbations, we analysed expression changes across all genes within 1 Mb upstream and downstream of the targeted loci. Most positive control perturbations (86%) resulted in at least one additional significantly differentially expressed gene (DEG). However, whilst *expected* genes were recovered with two or more independent gRNAs in most cases (83%) (Fig. 3A, S3A), over two thirds (68.3%) of all additional DEGs were statistically significant with a single gRNA (Fig. S3A). This lack of reproducibility between different gRNAs for the same target could be the result of variable on-target efficiencies, off-target effects and/or a sign of calibration issues of the statistical test. It is well recognised that differential gene expression testing using single-cell data results in the loss of appropriate false discovery rate (FDR) control when the lack of independence between cells from the same sample is not accounted for (Zimmerman et al. 2021). To assess whether this is the case for our data, we performed differential gene expression analysis for cells bearing non-targeting gRNAs, whose expression should not result in significant DEGs. Indeed, we observed that p-values were overly significant (Fig. S3B), resulting in an excess of positive calls.

To assess whether this is a problem specific to MAST, we tested two other differential gene expression algorithms, limma-voom (Law et al. 2014) and SCEPTRE (Barry et al. 2021), along with a non-parametric test (Wilcoxon rank sum test). All algorithms produced inflated p-values (Fig. S3C). When comparing the significant gRNA-DEG pairs reported by each method, we observed that a large fraction were only identified by one of the four methods (Fig. S3D); these corresponded almost entirely (94.2%) to genes other than the *expected* DEG, supporting that these are likely false positives. Instead, the *expected* genes from positive control perturbations were consistently detected by all four methods (Fig. S3D), building confidence in our experimental assay. Additionally, we observed that gRNA-DEG pairs reported by two or more methods were frequently supported by two or more independent gRNAs, while the method-specific pairs were most often identified with a single gRNA (Fig. S3E).

From all the methods considered, MAST showed the best recovery of *expected* gene expression changes along with the fewest method-specific pairs. Thus, we decided to use MAST results for all downstream analyses. To increase confidence in this set of DEGs, we exploited the evidence provided by independent gRNAs targeting the same element. We used Fisher’s method to integrate the p-values from all four gRNAs from each target into a single statistic – a *target-level p-value* – that reflects the amount of evidence supporting a gene expression change in perturbed cells (Fig. 4A). After adjusting for the multiple tests performed for the whole library, we obtained 378 significant DEGs (FDR < 5%; Table S2). We further classified these DEGs into confidence tiers, based on how many gRNAs supported the expression change (see Materials and Methods). Altogether, we identified 87 high-confidence expression changes supported by 3 or 4 gRNAs; 140 medium-confidence DEGs supported by 2 gRNAs; and 151 low-confidence effects supported by a single gRNA (Fig. 4A, Table S2). Since expression differences observed with a single gRNA are more likely to be off-target effects, we focused our analyses only on medium and high-confidence results.

**Figure 4:**
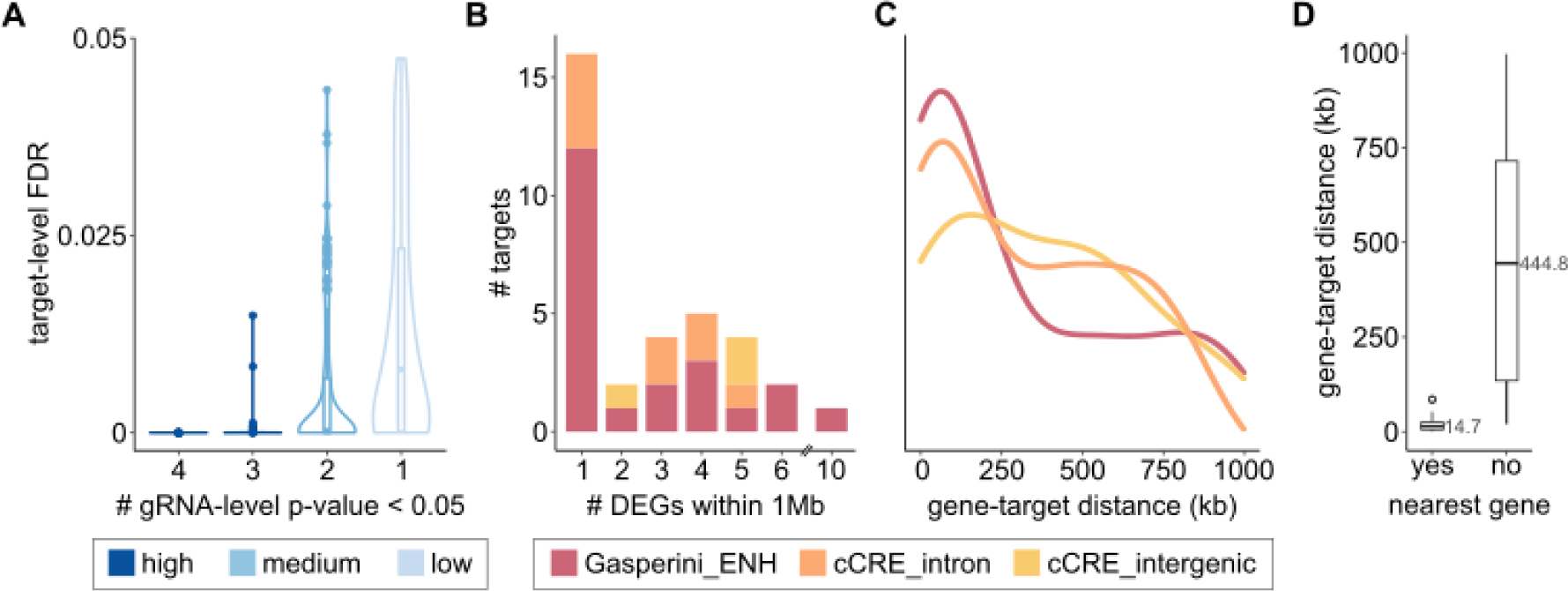
**A)** Distribution of the target-level adjustedp-values (FDR) for all significant element-to-gene (E2G) pairs detected, split by how many gRNAs have raw gRNA-level p-values < 0.05. E2G pairs supported by three or four gRNAs are high-confidence (dark blue); E2G pairs supported by 2 gRNAs are medium-confidence (blue); and E2G pairs supported by a single gRNA are low-confidence (light blue) and were discarded from downstream analyses. **B)** Barplot indicating the number of high and medium-confidence significant differentially expressed genes (DEGs) detected for non-coding perturbations within 1Mb up/downstream of the target site. Targets from Gasperini enhancers are shown in red; ENCODE cCREs are shown in orange if they lie within a gene intron, and in yellow if they are intergenic. **C)** Density plot of the distance between the E2G pairs, in kilobases. DEGs from targets of different classes are shown separately, as indicated by the same colours used in B. **D)** Boxplots of the distance between E2G pairs (as in C) but split by whether the gene is the nearest expressed gene to the target. The median is indicated.

For perturbations targeting Gasperini enhancers and cCREs (Fig. 2A) we detected 94 E2G pairs. These included at least one significant DEG for 22 of the 28 Gasperini perturbations; 9 of the 11 cCREs overlapping introns; and for all three cCREs in intergenic loci (Table S2). In general, non-coding perturbations did not induce widespread changes in expression. Almost half of all perturbations (47%) resulted in a single DEG within 1Mb of the target site, and an additional 32% resulted in fewer than five DEGs (Fig. 4B). These results suggest that CRISPRi effects are specific to the targeted elements. The majority (70.1%) of these non-coding element perturbations resulted in dysregulation of the nearest expressed gene, located a median distance of 15kb away from the target site (interquartile range (IQR): 5.5-25.8 kb; Fig. 4C-D). However, we also identified long-range effects affecting other genes, with a median distance of 445kb (IQR 135-715kb; Fig. 4C-D).

### Primary T cell crisprQTL helps interrogate disease-associated loci

Next, we focused on three case studies to emphasise how crisprQTL can aid in the identification of effector genes for disease GWAS associated regions. First, where expression data in the relevant cell type is publicly available, our crisprQTL data replicates colocalisations between disease GWAS signals and eQTL signals. For example, when perturbing the enhancer linked to *GIGYF1* by Gasperini *et al*. (2019) (Gasperini et al. 2019), which overlaps SNPs in the 95% credible set for a type 2 diabetes (T2D) GWAS signal (Fig. 5A) (Vujkovic et al. 2020), we found that *GIGYF1* was significantly differentially expressed (FDR=5.14e^-14^), with perturbed cells showing approximately a 10% reduction in expression (Fig. 5B). Furthermore, the T2D GWAS signal colocalised with an eQTL signal for *GIGYF1* in CD4^+^ T cells (Fig. 5C) (Chen et al. 2016, Javierre et al. 2016) and increased T2D risk colocalised with lower *GIGYF1* expression (Fig. 5D).

**Figure 5:**
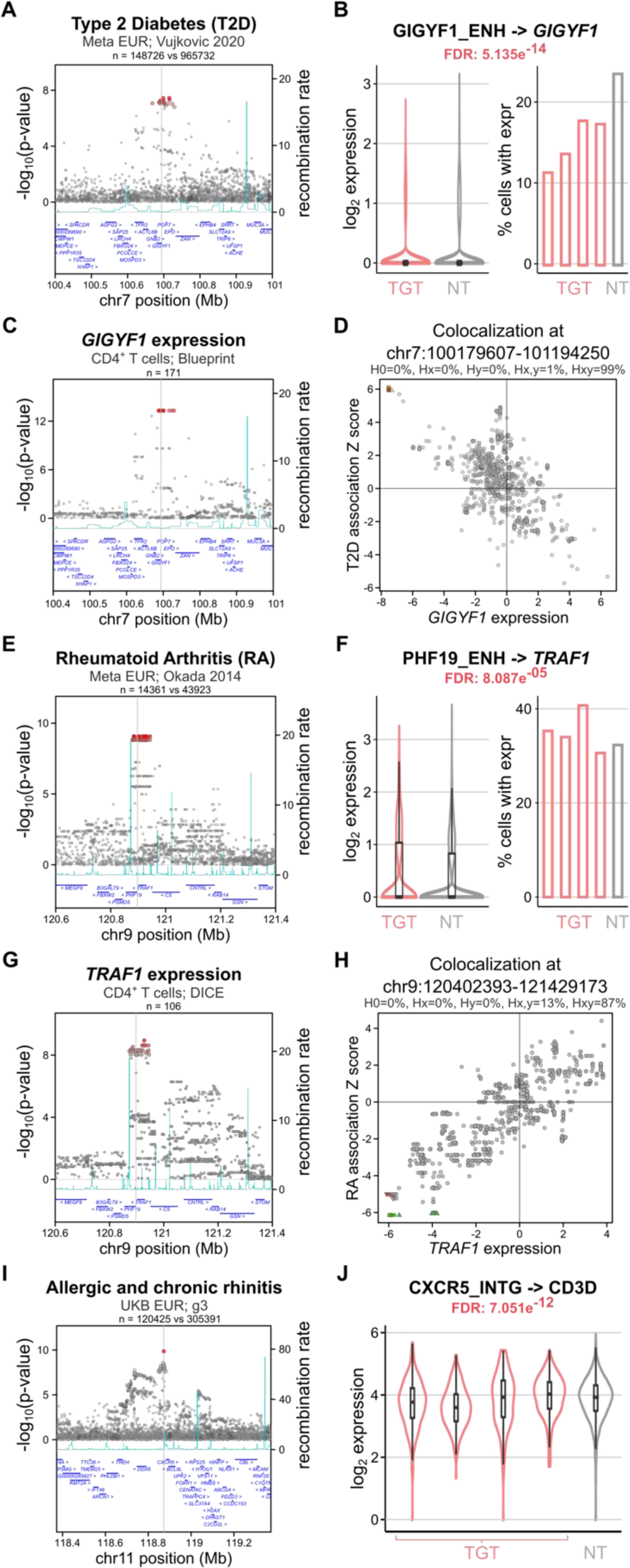
**A)** GWAS regional associationplot for type 2 diabetes (T2D) (Vujkovic, et al. 2020) highlighting the perturbed region with the grey line near *GIGYF1*. **B)** On the left, violin plots showing the normalised expression values of *GIGYF1* in cells expressing *GIGYF1* enhancer targeting (TGT) gRNAs (red) versus non-targeting (NT) controls (grey). On the right, barplot indicating the proportion of cells with TGT or NT gRNAs where expression of GIGYF1 is detected(counts>0). The target-level corrected p-value (FDR) of expression change is indicated at the top. **C)** eQTL regional association plot for *GIGYF1* expression in naïve CD4^+^ T cells (Chen et al. 2016, Javierre et al. 2016), highlighting the perturbed region in grey. **D)** Colocalisation plot of the T2D GWAS signal and *GIGYF1* eQTL in naïve CD4^+^ T cells, showing that these signals have a 99% posterior probability of being shared. T2D risk colocalises with decreased *GIGYF1* transcript expression. **E)** GWAS regional association plot for rheumatoid arthritis (RA) (Okada et al. 2014) highlighting the perturbed region in grey near *PHF9* and *TRAF1*. **F)** Same as D but for expression of *TRAF1* in cells expressing *PHF19* enhancer targeting gRNAs (red) versus NT controls (grey). **G)** eQTL regional association plot for *TRAF1* expression in naïve CD4^+^ T cells, highlighting the perturbed region with the grey line (Schmiedel et al. 2018, Chandra et al. 2021, Schmiedel et al. 2022). **H)** Colocalisation of the RA GWAS signal and *TRAF1* eQTL in naïve CD4^+^ T cells, showing that these signals have an 87% posterior probability of being shared. RA risk colocalises with increased *TRAF1* transcript expression. I) GWAS regional associationplot for allergic andchronic rhinitis (Bycroft et al. 2018) highlighting the perturbed area with a grey line near *CXCR5*. **J)** Violin plots showing the normalised expression values of *CD3D* in cells expressing targeting (TGT) gRNAs for *CXCR5* intergenic element (red) versus NT controls (grey). The target-level correctedp-value (FDR) of expression change is indicated at the top.

Second, for some enhancers, we identified E2G pairs in T cells that were different from those reported by Gasperini *et al*. (2019) in K562 cells (Gasperini et al. 2019), suggesting cell-type specific regulation. For instance, the perturbation of an enhancer element upstream of *PHF19* (Fig. 5E) that overlaps SNPs in the 95% credible set for a rheumatoid arthritis GWAS signal (Okada et al. 2014) resulted in immune-related DEGs not reported in K562 cells (Gasperini et al. 2019), including *TRAF1* (Fig. 5F). This rheumatoid arthritis GWAS signal colocalised with a *TRAF1* eQTL signal in CD4^+^ T cells (Schmiedel et al. 2018, Chandra et al. 2021) (Fig. 5G) and increased risk for rheumatoid arthritis colocalised with increased expression of *TRAF1* (Fig. 5H). This exemplifies how crisprQTL can help resolve cell-type specific putative effector genes at GWAS loci.

Finally, our method can also be used to identify novel E2G links that are missed with other approaches. For example, when perturbing an intergenic element upstream of *CXCR5* we detected differential expression of *CD3D,* whose TSS is more than 500 kb away from the perturbed region (Fig. 5I-J). The protein encoded by *CD3D* is a key component of the CD3 T cell co-receptor that plays essential roles in the adaptive immune response. The target perturbed region overlaps credible set SNPs for various immune-related traits, including asthma (Bycroft et al. 2018), multiple sclerosis (Patsopoulos et al. 2019), Fig. 5I), atopic dermatitis (Paternoster et al. 2015), rheumatoid arthritis (Okada et al. 2014), and rhinitis (Bycroft et al. 2018), and we found no eQTL colocalisation evidence supporting this E2G link. Together, our data suggest that crisprQTL can identify and validate cell-type specific E2G links supported by genetic evidence as well as discover novel associations that are missed in population-based studies.

## Discussion

We present a novel methodology called primary T cell crisprQTL – a CRISPRi-based single-cell functional screening approach in primary human CD4^+^ T cells that enables the mapping of regulatory elements to effector genes. Thus far, GWAS has identified a plethora of unique immune disease associations, many of which reside in cCREs in T cells (Trynka and Raychaudhuri 2013, Farh et al. 2015, Mumbach et al. 2017, Finucane et al. 2018, Calderon et al. 2019, Soskic et al. 2019, Nathan et al. 2021, Ota et al. 2021, Bossini-Castillo et al. 2022). Understanding the molecular mechanisms governed by these immune disease-associated regulatory elements would aid the understanding of disease aetiology and pave the way towards novel therapeutics (Nelson et al. 2015, King et al. 2019). The primary challenge, however, is identifying effector genes. Traditionally, eQTL studies have been used for this purpose however, eQTLs only clarify the effector gene at a limited fraction of GWAS signals (Chun et al. 2017, Yao et al. 2020, Umans et al. 2021, Connally et al. 2022, Mostafavi et al. 2022, Fletcher 2023). Furthermore, systematic studies have shown that GWAS and eQTL methods are biased to identify different types of variants, with eQTLs being depleted at functionally important variants (Mostafavi et al. 2022). Therefore, the use of functional methodologies, such as crisprQTL, provides a complementary approach for gene mapping.

We propose crisprQTL in disease-relevant cell models as a powerful functional genomics tool to help refine effector gene mapping at genetically-supported target loci. In this study we defined bona fide controls to enable future large-scale crisprQTL screens aiming to interrogate immune disease-associated elements in primary T cells. Beyond TSSs and locus control regions, we selected enhancer-element pairs previously identified in the K562 chronic myelogenous leukaemia cell line (Gasperini et al. 2019) and applied filters to ensure that: (1) the elements overlapped open chromatin in primary T cells, and (2) the gene was expressed in primary T cells. This approach can be applied for the development of crisprQTL studies in further disease relevant models. Out of the 28 prioritised enhancers, 17 mapped to the same effector gene in T cells and K562 cells, indicating conserved function. However, 12 showed distinct and/or additional gene targets, indicating cell-type specific regulation. For example, we perturbed an enhancer upstream of *PHF19* that was mapped to *PHF19* in crisprQTL studies in K562 cells (Gasperini et al. 2019) but to *TRAF1* in our primary T cell crisprQTL screen. Our observation was corroborated by colocalisation of a rheumatoid arthritis GWAS signal overlapping the enhancer with a *TRAF1* eQTL signal in CD4^+^ T cells (Fig. 5D-G). These findings highlight the importance of applying functional E2G mapping approaches in relevant cell models.

Recent advances in single-cell protocols, together with a drop in sequencing costs, are facilitating the generation of larger datasetsquerying cells derived from multiple donors, up to whole patient cohorts. The inclusion of higher biological replication will limit false discoveries and improve power to detect subtle perturbation effects (Squair et al. 2021). Furthermore, the single-cell resolution inherent to crisprQTL enables the study of the impact of cell subtypes andstates on the complex interrelationships between genetic variation, regulatory elements and genes. This will provide a valuable resource for higher throughput crisprQTL screens in primary, dynamic and complex cell models and will enable delineating cell subtype- or state-specific regulation driven by disease-associated variants (Mumbach et al. 2017, Zhernakova et al. 2017, Schmiedel et al. 2018, van der Wijst et al. 2018, Donovan et al. 2020, Kim-Hellmuth et al. 2020, Chiou et al. 2021, Ota et al. 2021, Young et al. 2021, Nathan et al. 2022, Soskic et al. 2022, Yazar et al. 2022). For example, studies modelling eQTLs obtained from single-cell RNA-seq data from primary T cells helped disentangle cell subtype-specific effects that were masked in bulk RNA-seq(Soskic et al. 2022). Others have identified loci with independent eQTLs that have opposing state-dependent effects (Nathan et al. 2022). Furthermore, using H3K27ac HiChIP data in primary human naïve T cells, regulatory T cells (Treg) and T helper 17 cells (Th17), Mumbach, *et al*. (2017) identified links between autoimmune disease variants and effector genes that were present in T cell effector types (Treg and Th17) but not in naïve T cells (Mumbach et al. 2017).

To enable the perturbation of regulatory elements, we established CRISPRi in primary CD4^+^ T cells by lentiviral transduction of both dCas9-repressor and a pooled gRNA library. A similar approach (Schmidt et al. 2022) was used for gene silencing in primary T cells using the KOX1 KRAB domain, to enable fluorescence-activated cell sorting-based screens. However, we show that the ZIM3 domain is a more potent repressor in primary T cells (Fig. 1D), consistent with previous observations in cell lines (Alerasool et al. 2020). This improved perturbation efficiency is important when targeting non-coding regulatory elements, since this generally results in much smaller effects on gene expression compared to TSS silencing. While we and others have shown that pooled CRISPRi screens coupled to scRNA-seq enable unbiased interrogation of hundreds of elements in a single assay (Gasperini et al. 2019, Morris et al. 2023), other perturbation methods for the discovery and mechanistic study of E2G associations are rapidly evolving and could provide orthogonal means of validation. For instance, CRISPR/Cas9 directed mutagenesis has been employed to interrogate non-coding regulatory elements at defined loci (Canver et al. 2015, Diao et al. 2016, Fulco et al. 2016, Korkmaz et al. 2016, Rajagopal et al. 2016, Sanjana et al. 2016, Gasperini et al. 2017). Although these methods suffer from limited scalability, their application to prioritised loci could uncover regulatory maps at much higher precision. CRISPRa and other CRISPR epigenetic editing methods have also been employed to study non-coding regulation in small-scale studies (Amabile et al. 2016, Klann et al. 2017, Li et al. 2020). Moving forward, a systematic comparison and benchmarking of E2G perturbation maps generated with multiple CRISPR modifiers and precise edits across multiple tissues should help us define robust principles for E2G interrogation and discovery.

In summary, we present an experimental perturbation approach and analytical pipeline for the validation and discovery of E2G links in primary human T cells. Our method and the data presented here set the basis for future functional interrogation of immune disease-associated distal elements at larger scale and in other *in vitro* primary cell models. Together with population-based studies, crisprQTL E2G maps across multiple human tissues will aid our understanding of the effector genes and pathways driving human disease.

## Materials and Methods

### Locus selection

The genomic coordinates for the three *CD2* LCR were obtained from (Lake et al. 1990, Festenstein et al. 1996, Kaptein et al. 1998).

We selected enhancers from the set of 470 high-confidence intergenic E2G pairs from Gasperini *et al*. (2019) with the following features: 1) the putative enhancer must overlap a region of open chromatin in primary CD4^+^ T cells as identified in at least one of the following datasets: i) ATAC-seq peaks identified on CD4^+^ T naïve or memory cells after 16 hours or 5 days of CD3/CD28 activated stimulation (Soskic et al. 2019) or ii) ATAC-seq peaks from multiple CD4^+^ T cells from (Calderon et al. 2019); 2) the *expected* target gene, as defined by Gasperini *et al*. (Gasperini et al. 2019), was amongst the top 50% expressed genes in primary T cells, as assessed from a scRNA-seqdataset from CD3/CD28 stimulated CD4^+^ T cells (Cano-Gamez et al. 2020). After applying these filters, a set of 28 E2G pairs were selected.

Additionally, we selected 11 intronic and 3 intergenic cCREs (ENCODE-Project-Consortium 2012) overlapping open chromatin regions in primary T cells (Calderon et al. 2019).

### gRNA design

Four gRNAs were included per target in the pooled library. gRNAs targeting TSSs were selected from the Dolcetto library (Sanson et al. 2018). gRNAs targeting non-coding elements (*CD2* LCRs, enhancers selected from Gasperini *et al*. (2019) (Gasperini et al. 2019), and ENCODE cCREs), were designed using the Broad Institute platform <u>CRISPick</u> (Sanson et al. 2018) and selected to target the middle 300bp region of the element (Table S1). For enhancers selected from Gasperini *et al*. (2019), two out of the four gRNAs overlapped the gRNAs used in the original study by Gasperini *et al*. (2019) (Gasperini et al. 2019). Lastly, 35 non-targeting gRNAs were included in the pooled library, selected from the Dolcetto library (Sanson et al. 2018). gRNA sequences for each element are reported in Table S1.

### gRNA cloning and pooled gRNA library construction

Individual gRNAs and the pooled library were cloned into a CROPseq lentiviral backbone adapted from (Datlinger et al. 2017) with the following modifications: 1) the scaffold sequence was modified to include the optimized CRISPRi scaffold described in (Chen et al. 2013); and 2) a fluorescent ZsGreen marker was included downstream of the EF1α promoter and linked through T2A to a puromycin resistance cassette. This backbone is referred to as CRISPRi CROPseq.

#### Cloning of individual gRNAs

To clone gRNAs into this backbone (Genewiz), oligos containing the protospacer sequence and recombination arms homologous to the vector backbone were synthesised (oligo structure: 5’ tttcttggctttatatatcttgtggaaaggacgaaacacc-protospacer-gtttaagagctatgctggaaacagcatagcaagtttaaat 3’). The CRISPRi-seq backbone was digested with AarI and then cloned with the insert by recombination-based cloning. The reaction mixture was transformed into DH5α competent cells and the colonies were selected at 37°C. Individual colonies were picked and plasmid DNA was isolated and verified by Sanger sequencing. The sequences of TSS-targeting gRNAs and controls used in Fig. 1 and Fig. S1 are detailed in the following table:

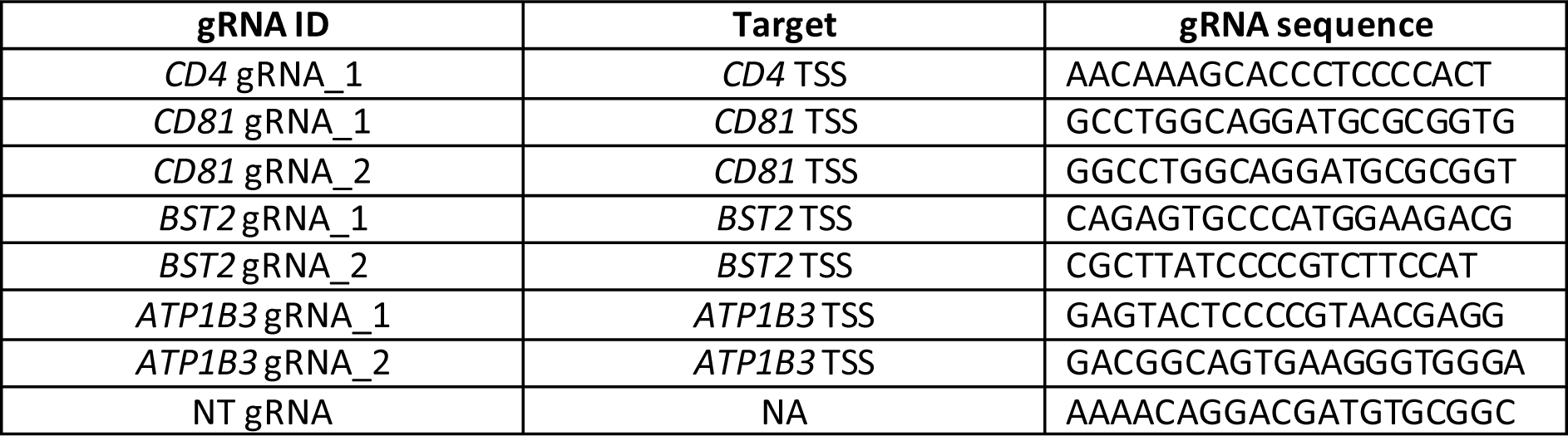

#### Cloning of pooled library (355 gRNAs)

gRNA oligos were synthesized, PCR amplified and cloned into the CRISPRi CROPseq backbone using Gibson assembly, as previously described (Joung et al. 2017), by VectorBuilder. The pooled gRNA library was built from a total of 7×10^5^ single colonies, which represents more than 1000-fold coverage of the designed gRNAs. Library gRNA representation was assessed by 150 bp paired-end sequencing (Illumina, NovaSeq). The distribution of gRNA representation is shown in Fig. S2D.

### Lentiviral vector production and determination of viral titre

HEK293T suspension adapted cells (in-house) were cultured in growth media consisting of BalanCD HEK293 medium (Irvine Scientific, 91165) supplemented with 2% GlutaMAX (Thermo Fisher Scientific, 35050061) and 1% Pluronic F-68 (Thermo Fisher Scientific, 24040032). For lentiviral packaging, cells were seeded at 2×10^6^ cells/ml density and transfected with a total 190 ug DNA complexed with 450 ug PEIpro (Polyplus-transfection, 101000033). The plasmid DNA mix, which included one envelope encoding *VSVg*, two packaging plasmids encoding *rev*, and *gag* and *pol* genes, and a transfer DNA plasmid vector, was added to pre-warmed OptiMEM (Thermo Fisher Scientific, 31985070), followed by addition of PEIpro and incubation for 30 min at room temperature for the transfection complexes to form. The transfection mixture was added to the HEK293T suspension adapted cells and the cells were incubated in a shaker cell culture incubator (Multitron shaker incubator, Infors HT) at 37°C with 5% CO_2_ at 110 rpm. The next day, 1.5mM sodium butyrate (Sigma, 303410) was added to the transfected cell culture. Lentiviral vector-containing media from the transfected cells was collected 72h after transfection, clarified by centrifugation at 500 *x g* for 10 min, filtered through 0.45 μm filter units and concentrated via high-speed centrifugation at 70,000 *x g* for 2h. The viral pellet was then resuspended in RPMI media (Gibco, 11534446) at a ∼350X concentration. The resuspended lentiviral vector solution was aliquoted and stored at −80°C.

Functional viral titre was determined by transducing 1×10^5^ HEK293T adherent cells (Lenti-X™ 293T Cell Line, 632180) with serial dilutions of the lentiviral solution in a total volume of 100 μL per well of a 96-well plate and incubated at 37°C in 5% CO_2_. For lentiviral vectors containing the fluorescent marker ZsGreen, the percentage of cells expressing ZsGreen was quantified three days post-transduction by flow cytometry using Cytoflex S (Beckman) flow cytometer, normalised to the starting cell number and dilution factor to obtain the titre in transduction units per mL (TU/mL).

### Isolation and culture of primary human CD4^+^ T cells

The human biological samples in this study were sourced ethically and their research use was in accord with the terms of the informed consent under an IRB/EC approved protocol. Briefly, mononuclear cells from circulating blood were removed by apheresis. From these leukopaks, primary human CD4^+^ T cells were enriched using CliniMACS Prodigy setup and human CD4 MicroBeads (Miltenyi, 130-045-101). Following positive selection, cells were aliquoted and cryopreserved. Primary CD4^+^ T cell vials were thawed at the time of the experiments and cells cultured in RPMI 1640 (Gibco, 11340892) supplemented with 10% heat-inactivated FBS, 1x Glutamax (Thermo Fisher Scientific, 35050061), 1mM Sodium Pyruvate (Thermo Fisher Scientific, 11360088), 5mM HEPES (Thermo Fisher Scientific, 15630080), 1x non-essential aminoacids (Thermo Fisher Scientific, 11140035), 1x Penicillin/streptomycin (Thermo Fisher Scientific, 15070063), 55 µM 2-mercaptoethanol (Thermo Fisher Scientific, 21985023), and 15 ng/mL of recombinant human interleukin 2 (IL-2) (Peprotech, 200-02). Cells were kept in a humidified 5% CO_2_ atmosphere at 37°C.

### dCas9-KRAB transduction and selection

Primary human CD4^+^ T cells were thawed and cultured as described above at a density of ∼1×10^6^ cells per mL. The next day, T cells were activated using Dynabeads Human T-Activator CD3/CD28 (Thermo Fisher Scientific, 11131D), at 1:1 cell:bead ratio, according to manufacturer’s guidelines. 16-20h after activation, cells were transduced with the corresponding lentiviral vector encoding a KRAB-dCas9 fusion under different promoters, and a blasticidin resistance gene. Lentiviral transductions were performed in growth media supplemented with 5mM HEPES (Thermo Fisher Scientific, 15630080), and spinoculation was carried out at 800 *x g* for 1h at 37°C. 72h after transduction, blasticidin selection was carried out at 20 ug/mL for two days, followed by an additional 5 days at 12.5 ug/mL. Media was replenished and cells were expanded as necessary based on confluency.

KRAB-dCas9 lentiviral constructs tested in this study include: SFFV-dCas9-KOX1-T2A-Bsd, EFS-ZIM3-dCas9-P2A-Bsd, EFS-KOX1-dCas9-P2A-Bsd, CBh-ZIM3-dCas9-P2A-Bsd, CBh-KOX1-dCas9-P2A-Bsd and SFFV-dCas9-KOX1-MeCP2-T2A-Bsd (Fig. 1C). Blasticidin selection of cells transduced with SFFV-dCas9-KOX1-MeCP2-T2A-Bsd resulted in no to very little viable cells, reflecting inefficient lentiviral transduction, likely due to the large provirus size.

### gRNA transduction and flow cytometry analysis of CRISPRi efficiency

Following one day of rest after blasticidin selection, KRAB-dCas9 expressing cells were activated with Dynabeads Human T-Activator CD3/CD28 (Thermo Fisher Scientific, 11131D), at 1:1 cell:bead ratio. 16-20h after activation, cells were transduced with a CRISPRi CROPseq lentivirus encoding gRNAs targeting either *CD4*, *CD81*, *BST2* or *ATP1B3*, or a non-targeting control (NT) gRNA, at a MOI of 0.1, in growth media supplemented with 5mM of HEPES (Thermo Fisher Scientific, 15630080). Spinoculation was carried out at 800 *x g* for 1h at 37°C. 48h after gRNA transduction, cells were selected with 2 ug/mL puromycin for 4 days, and selection was verified by ZsGreen expression in >95% of the cells by flow cytometry (Cytoflex S, Beckman Coulter).

Flow cytometry was performed at different timepoints to estimate CRISPRi efficiency by measuring protein downregulation of the target genes (*CD4*, *CD81*, *BST2* and *ATP1B3*) compared to a NT gRNA. Briefly, ∼100,000 cells were washed with PBS and stained for 1h at 4°C with an antibody targeting the corresponding protein. Next, cells were washed with PBS and analysed by flow cytometer (Cytoflex S, Beckman Coulter), recording 30,000-50,000 events. Antibodies used were APC anti-human *CD4* clone RPA-T4 (Biolegend, 300514), APC anti-human *CD81* clone 5A6 (Biolegend, 349510), APC anti-human *CD317* (BST2, Tetherin) clone RS38E (Biolegend, 348410), and APC-anti-human *CD298* clone LNH-94 (Biolegend, 341706).

### Pooled CRISPRi screen

Primary human CD4^+^ T cells expressing CBh-ZIM3-dCas9-Bsd (transduced and selected as described above) were activated with Dynabeads Human T-Activator CD3/CD28 (Thermo Fisher Scientific, 11131D) at 1:1 cell:bead ratio and, 16-20h later, two million cells were transduced with the pooled CRISPRi CROPseq gRNA library (described above) at a <0.3 MOI. The transduction was performed in growth media supplemented with 5mM of HEPES (Thermo Fisher Scientific, 15630080) and spinoculation was carried out at 800 *x g* for 1h at 37°C. 48h later, cells were selected with 2 ug/mL puromycin for 4 days, and selection was verified by ZsGreen expression in >95% of the cells by flow cytometry (Cytoflex S, Beckman Coulter). Cells were cultured in growth media for an additional 6 days before processing for scRNA-seq.

### Preparation of 10X Genomics scRNA-seq libraries and sequencing

The pooled CRISPRi CROPseq screen was read-out using the 10X Genomics platform (10X Genomics, PN-1000075). The screen dataset contains two different scRNA-seq runs: the first one (8 channels of a 10X Genomics chip B, PN-1000073) was performed with fresh cells at the end of the CRISPRi experiment, 10 days post gRNA library transduction; the second one (32 channels across four 10XGenomics chips B, PN-1000073) was performed using cells from the same experiment that were cryopreserved 10 days after gRNA transduction and thawed in puromycin-containing media (2 ug/ml) three days before the 10X Genomics run. To remove dead cells and debris, the cell suspension was treated with Lymphoprep (Axis-Shield, 12HHS09) the day before loading the cells into the 10X Chromium Controller. Each channel from a 10X Genomics chip B (PN-1000073) was loaded with 16,000 cells into a Chromium Controller, following manufacturer’s guidelines. 3’ gene expression libraries (v3) were prepared following manufacturer’s instructions (10X Genomics, PN-1000075). gRNA amplicon libraries were generated from amplified cDNA following the protocol described in (Alda-Catalinas et al. 2021). Gene expression and gRNA libraries were QCed by tapestation (Agilent) and quantified by Qubit (Thermo Fisher, Q32851) and KAPA qPCR (Kapa Biosystems, KK4824). Multiplexed gene expression libraries were pooled at a 5:1 ratio with multiplexed gRNA libraries. The pool of gene expression and gRNA libraries were sequenced across two S4 and one S2 flow cells of a Novaseq 6000 with 28 cycles for read 1, 91 cycles for read 2 and 8 cycles for i7 index.

### CROP-seq data processing

Sequencing data were processed using cellranger count v4.0, with default parameters. We used the human reference genome provided by 10X Genomics (hg38, Ensembl annotation version 98; https://cf.10xgenomics.com/supp/cell-exp/refdata-gex-GRCh38-2020-A.tar.gz), supplemented with artificial chromosomes containing the sequences for all gRNAs present in the library to enable recovery of gRNA-derived transcripts from the cDNA library. Additionally, a feature reference file containing all gRNA sequences present in the library was supplied to quantify UMI counts for each gRNA in the gRNA library. The barcode-rank plots for all samples were concordant with good-quality data and appropriate distinction of cell-containing from empty droplets. A median of 7,218 cells (s.d. 1445 cells) were recovered from each technical replicate. For downstream analyses, the filtered count matrices produced by cellranger were imported into R using the dropletUtils package (Griffiths et al. 2018, Lun et al. 2019).

### Quality control

To remove cells of poor quality we used scater (McCarthy et al. 2017) to compute QC metrics. For each barcode, we assessed the total UMI counts, total number of genes detected, and the proportion of counts mapping to mitochondrial genes (Fig. S2A). We removed any barcodes that deviated by more than 3 median absolute deviations (MAD) from the median for any of the three metrics’ distributions. To account for slight differences in sequencing depth between 10X chips, thresholds were defined independently for each sample (Fig. S2A). Overall, we removed 4.07% of the barcodes. We further identified outlier barcodes with very small fraction of mitochondrial reads (deviating by more than 3 MADs), as these are likely to correspond to stripped nuclei, instead of cells. We confirmed that these barcodes had a substantially lower number of genes detected and were thus removed from downstream analyses.

### Doublet detection

To identify putative doublets, we used the method implemented in the scDblFinder package (Germain et al. 2021) with default parameters. Barcodes identified as doublets had higher total UMI counts, number of detected genes, and were more likely to have more than one gRNA, and were thus discarded. A total of 250,195 barcodes were retained, representing singlet, good-quality cells.

### gRNA assignment

To determine which gRNAs were present in each cell we applied a binomial test to the UMI counts obtained from the gRNA library (see above). The probability of success of each gRNA was determined from the initial representation of each gRNA in the lentiviral library, assessed by DNA sequencing. The binomial test takes into account the total library size of each cell, together with the expected proportions of each gRNA, to determine the minimum UMI count required to consider a gRNA present above background noise levels. Any gRNAs with a Bonferroni-adjusted p-value smaller than 0.001 were considered present in that cell. We additionally discarded any significant gRNA assignments that were supported by 3 UMI counts or fewer. Overall, we were able to confidently assign gRNA calls to 152,403 cells (60.91%). From these, almost all (95.62%) had a single gRNA assigned, consistent with experiments carried out at low multiplicity of infection.

### Normalisation and batch correction

Gene expression counts were normalised using the deconvolution method implemented in scran (Lun et al. 2016). Highly variable genes were inferred with the modelGeneVar function, and the top 2000 most variable genes were used for dimensionality reduction (PCA followed by UMAP, as implemented in scater). Data visualisation indicated strong separation of the two experiments performed. Thus, the experiment was included as a covariate in all differential expression analyses.

### Differential expression analysis

To determine the effects from each perturbation, we used MAST (Finak et al. 2015) to test for gene expression differences between all cells containing a particular gRNA, compared to a set of randomly selected 5,000 cells containing only a non-targeting gRNA (referred to as NT cells). We restricted the analysis to genes that were detected in at least 5% of cells (10,047 genes). The model fit was done using all cells containing any of the four gRNAs for the same target, plus the background set of NT cells. We added as covariates the experiment of each sample and the detection rate as defined by MAST’s authors. Then, the fitted model was used to test the effect of each gRNA, by specifying the corresponding contrast. Results were filtered to include only genes that fall within 1Mb up/downstream of the target (defined from Ensembl’s v98 annotation, with the BiomaRt package (Durinck et al. 2005, Durinck et al. 2009)). P-values were adjusted for multiple testing with the Benjamin-Hochberg method.

To assess p-value calibration, we followed the same strategy as Barry *et al*. 2021 (Barry et al. 2021). Briefly, the 35 non-targeting (NT) gRNAs were grouped into nine groups of four, to emulate the library structure of targeting perturbations. MAST was run in the same way as described above. Since NT gRNAs do not have a genomic location, we considered all genes tested against any targeting gRNA (i.e. any gene within 1Mb of a targeted locus). The p-values reported by MAST were compared to expected p-values under the null (Fig. S3A).

The same strategy was used to test for changes in expression for both targeting and non-targeting gRNAs using limma-voom (Law et al. 2014) or SCEPTRE (Barry et al. 2021). For the Wilcoxon rank sum test, we used the implementation from the pairwiseWilcox function from the scran package (Lun et al. 2016), blocking on the experiment of each sample to account for the observed batch effect.

### Target-level perturbation effects

To integrate the results from the four gRNAs targeting the same genomic locus, we used Fisher’s method (as implemented in the combineGroupedPValues function from the metapod package (Lun 2022)) to integrate the raw p-values from all four gRNAs into a single p-value. These target-level p-values were corrected for multiple testing to account for all the tests performed for the complete library (considering genes within 1Mb of the target, accounting for 1,336 tests), using the Benjamini-Hochberg method. Genes with an adjusted target-level p-value < 0.05 were considered significantly differentially expressed (DEGs). DEGs were assigned to confidence tiers depending on the number of gRNAs with a raw p-value < 0.05 with concordant effects. High-confidence tier: 3 or 4 gRNAs; medium-confidence tier: 2 gRNAs; and low-confidence tier: 1 gRNA. Table S2 includes all significant E2G links identified by MAST analysis.

### GWAS credible set intersections

For a wide range of publicly available GWAS, we generated conditional test statistics (Yang et al. 2011, Yang et al. 2012) and conditional 95% credible sets from GWAS summary statistics using an established Bayesian fine-mapping approach (Wakefield 2009, Maller et al. 2012). For each GWAS signal which overlapped the perturbed target region +/− 1-kb we also checked for colocalisation of eQTL signals for the DEGs using the coloc software with default parameters (Giambartolomei et al. 2014, Wallace 2022).

## Data availability

The raw and processed data from this study will be available in a public repository (submission in progress). All code used for data processing and analysis will be released with the published manuscript.

## Acknowledgements

We would like to thank Sebastian Ullrich, Valeriia Sherina and Bin Sun for insightful discussions. This study makes use of data generated by the Blueprint Consortium. A full list of the investigators who contributed to the generation of the data is available from www.blueprint-epigenome.eu. Funding for the project was provided by the European Union’s Seventh Framework Programme (FP7/2007-2013) under grant agreement no 282510 BLUEPRINT”. This research has been conducted using data from UK Biobank www.ukbiobank.ac.uk, a major biomedical database.

## Supplemental Figures

**Figure S1:**
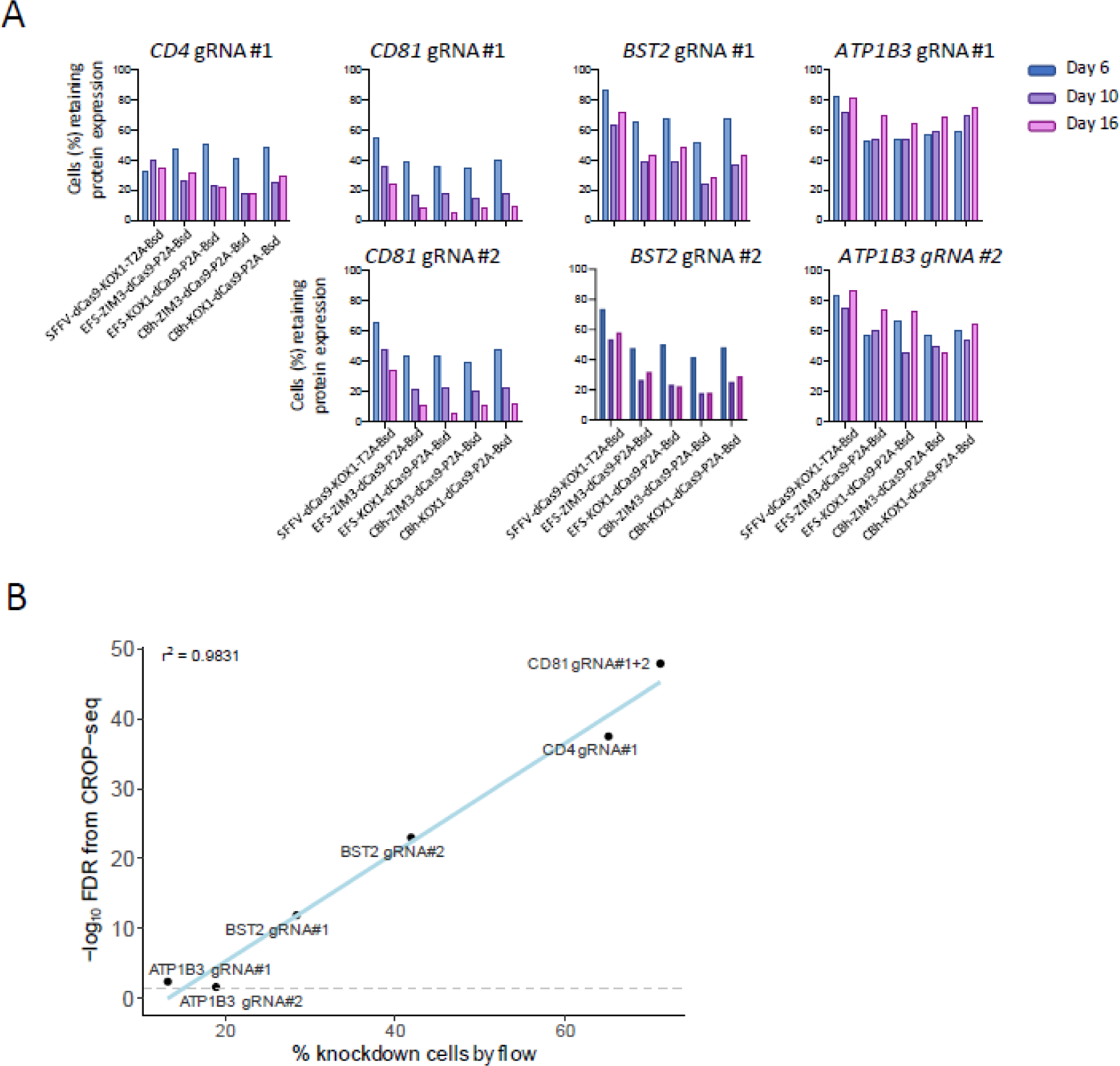
**A)** Bar plots showing percentage of cells retaining proteinexpression for different target genes (*CD4, CD81, BST2, ATP1B3*) and gRNAs at days 6, 10 and 16 post-gRNA transduction into primary CD4^+^ T cells expressing different dCas9-repressor constructs, analysed by flow cytometry and normalised to the corresponding NT gRNA control sample. **B)** Percentage of cells showing protein downregulation of the target gene after CRISPRi by flow cytometry (x-axis) versus significance (−log10 FDR) of downregulation of the target gene (mRNA) by 3’ scRNA-seq analysis (y-axis), normalised to the corresponding NT control. Pearson r^2^ = 0.98.

**Figure S2:**
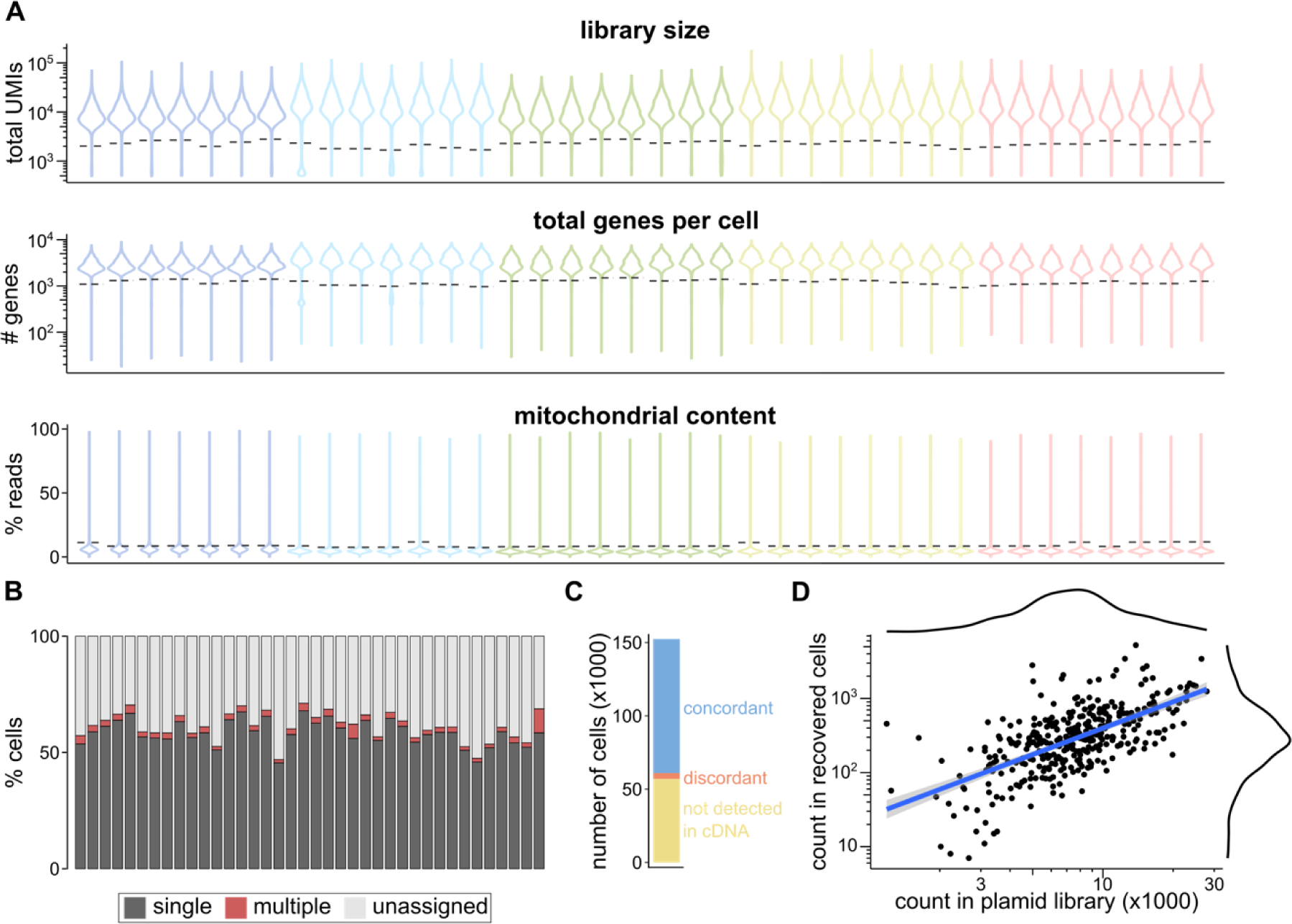
**A)** Distributions of the total UMIs per cell, number of detected genes per cell, and fraction of reads mapping to mitochondrial genes, used for quality control of the scRNAseq data. Each violin corresponds to a technical replicate (channel in a 10X chip); colours indicate different 10X chips. The dotted lines indicate the thresholds used for each replicate to exclude poor-quality cells. **B)** Same as Fig. 2C but split per technical replicate. **C)** Barplot showing the number of cells where the same gRNA is the most abundant in both the cDNA and gRNA libraries (concordant, blue); the most abundant gRNA is different between libraries (discordant, orange); or no gRNA information was recovered from the cDNA library (yellow) **D)** Scatter plot of the relative abundance of each gRNA in the plasmid library (*x-axis*, assessed by DNA sequencing of the library) versus the number of cells positive for each gRNA (*y-axis*, assessed from the scRNAseq data).

**Figure S3:**
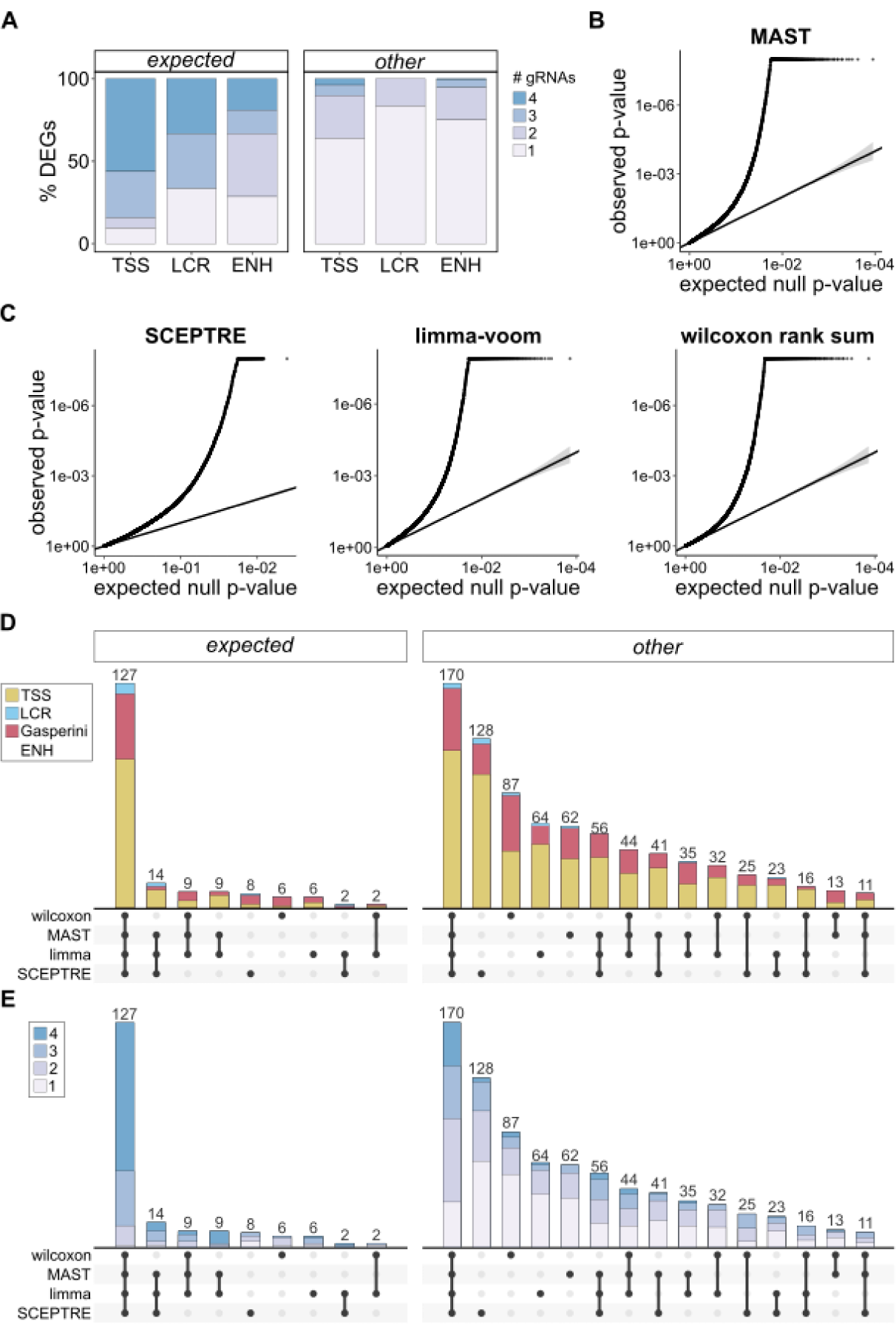
**A)** Barplots of the fraction of significant differentially expressed genes (DEGs) from positive control perturbations that are supported by different numbers of gRNAs (raw gRNA-levelp-value < 0.05). The *expected* genes are shown separately from all *other* DEGs. **B)** QQ plot of the expected vs observed p-values reported by MAST when testing for expression changes from non-targeting gRNAs, which should not induce any significant changes in expression. Deviation from the diagonal indicates inflated p-values. **C)** Same as B) but for results from SCEPTRE, limma-voom or a Wilcoxon rank sum test. **D)** Upset plot indicating the number of DEGs for positive controltargets that are identified byany of the four methods. The height of the bar indicates the number of DEGs, split by target class (indicated by different colours). Under each bar, the methods that called the gene as significant are indicated. The *expected* genes are shown separately from all *other* DEGs. **E)**Same as D but with the colours indicatingthe number of gRNAs supporting each DEG (raw gRNA-level p-value < 0.05).

## References

Alda-Catalinas, C., D. Bredikhin, I. Hernando-Herraez, F. Santos, O. Kubinyecz, M. A. Eckersley-Maslin, O. Stegle and W. Reik (2020). “A Single-Cell Transcriptomics CRISPR-Activation Screen Identifies Epigenetic Regulators of the Zygotic Genome Activation Program.” Cell Syst 11(1): 25–41.e29.

Alda-Catalinas, C., M. A. Eckersley-Maslin and W. Reik (2021). “Pooled CRISPR-activation screening coupled with single-cell RNA-seq in mouse embryonic stem cells.” STAR Protoc 2(2): 100426.

Alerasool, N., D. Segal, H. Lee and M. Taipale (2020). “An efficient KRAB domain for CRISPRi applications in human cells.” Nat Methods 17(11): 1093–1096.

Alsheikh, A. J., S. Wollenhaupt, E. A. King, J. Reeb, S. Ghosh, L. R. Stolzenburg, S. Tamim, J. Lazar, J. W. Davis and H. J. Jacob (2022). “The landscape of GWAS validation; systematic review identifying 309 validated non-coding variants across 130 human diseases.” BMC Med Genomics 15(1): 74.

Amabile, A., A. Migliara, P. Capasso, M. Biffi, D. Cittaro, L. Naldini and A. Lombardo (2016). “Inheritable Silencing of Endogenous Genes by Hit-and-Run Targeted Epigenetic Editing.” Cell 167(1): 219–232.e214.

Amariuta, T., Y. Luo, R. Knevel, Y. Okada and S. Raychaudhuri (2020). “Advances in genetics toward identifying pathogenic cell states of rheumatoid arthritis.” Immunol Rev 294(1): 188–204.

Barry, T., X. Wang, J. A. Morris, K. Roeder and E. Katsevich (2021). “SCEPTRE improves calibration and sensitivity in single-cell CRISPR screen analysis.” Genome Biol 22(1): 344.

Boix, C. A., B. T. James, Y. P. Park, W. Meuleman and M. Kellis (2021). “Regulatory genomic circuitry of human disease loci by integrative epigenomics.” Nature 590(7845): 300–307.

Bossini-Castillo, L., D. A. Glinos, N. Kunowska, G. Golda, A. A. Lamikanra, M. Spitzer, B. Soskic, E. Cano-Gamez, D. J. Smyth, C. Cattermole, K. Alasoo, A. Mann, K. Kundu, A. Lorenc, N. Soranzo, I. Dunham, D. J. Roberts and G. Trynka (2022). “Immune disease variants modulate gene expression in regulatory CD4(+) T cells.” Cell Genom 2(4): None.

Buniello, A., J. A. L. MacArthur, M. Cerezo, L. W. Harris, J. Hayhurst, C. Malangone, A. McMahon, J. Morales, E. Mountjoy, E. Sollis, D. Suveges, O. Vrousgou, P. L. Whetzel, R. Amode, J. A. Guillen, H. S. Riat, S. J. Trevanion, P. Hall, H. Junkins, P. Flicek, T. Burdett, L. A. Hindorff, F. Cunningham and H. Parkinson (2019). “The NHGRI-EBI GWAS Catalog of published genome-wide association studies, targeted arrays and summary statistics 2019.” Nucleic Acids Res 47(D1): D1005–d1012.

Bycroft, C., C. Freeman, D. Petkova, G. Band, L. T. Elliott, K. Sharp, A. Motyer, D. Vukcevic, O. Delaneau, J. O’Connell, A. Cortes, S. Welsh, A. Young, M. Effingham, G. McVean, S. Leslie, N. Allen, P. Donnelly and J. Marchini (2018). “The UK Biobank resource with deep phenotyping and genomic data.” Nature 562(7726): 203–209.

Calderon, D., M. L. T. Nguyen, A. Mezger, A. Kathiria, F. Müller, V. Nguyen, N. Lescano, B. Wu, J. Trombetta, J. V. Ribado, D. A. Knowles, Z. Gao, F. Blaeschke, A. V. Parent, T. D. Burt, M. S. Anderson, L. A. Criswell, W. J. Greenleaf, A. Marson and J. K. Pritchard (2019). “Landscape of stimulation-responsive chromatin across diverse human immune cells.” Nat Genet 51(10): 1494–1505.

Cano-Gamez, E., B. Soskic, T. I. Roumeliotis, E. So, D. J. Smyth, M. Baldrighi, D. Willé, N. Nakic, J. Esparza-Gordillo, C. G. C. Larminie, P. G. Bronson, D. F. Tough, W. C. Rowan, J. S. Choudhary and G. Trynka (2020). “Single-cell transcriptomics identifies an effectorness gradient shaping the response of CD4(+) T cells to cytokines.” Nat Commun 11(1): 1801.

Canver, M. C., E. C. Smith, F. Sher, L. Pinello, N. E. Sanjana, O. Shalem, D. D. Chen, P. G. Schupp, D. S. Vinjamur, S. P. Garcia, S. Luc, R. Kurita, Y. Nakamura, Y. Fujiwara, T. Maeda, G. C. Yuan, F. Zhang, S. H. Orkin and D. E. Bauer (2015). “BCL11A enhancer dissection by Cas9-mediated in situ saturating mutagenesis.” Nature 527(7577): 192–197.

Chandra, V., S. Bhattacharyya, B. J. Schmiedel, A. Madrigal, C. Gonzalez-Colin, S. Fotsing, A. Crinklaw, G. Seumois, P. Mohammadi, M. Kronenberg, B. Peters, F. Ay and P. Vijayanand (2021). “Promoter-interacting expression quantitative trait loci are enriched for functional genetic variants.” Nat Genet 53(1): 110–119.

Chen, B., L. A. Gilbert, B. A. Cimini, J. Schnitzbauer, W. Zhang, G. W. Li, J. Park, E. H. Blackburn, J. S. Weissman, L. S. Qi and B. Huang (2013). “Dynamic imaging of genomic loci in living human cells by an optimized CRISPR/Cas system.” Cell 155(7): 1479–1491.

Chen, L., B. Ge, F. P. Casale, L. Vasquez, T. Kwan, D. Garrido-Martín, S. Watt, Y. Yan, K. Kundu, S. Ecker, A. Datta, D. Richardson, F. Burden, D. Mead, A. L. Mann, J. M. Fernandez, S. Rowlston, S. P. Wilder, S. Farrow, X. Shao, J. J. Lambourne, A. Redensek, C. A. Albers, V. Amstislavskiy, S. Ashford, K. Berentsen, L. Bomba, G. Bourque, D. Bujold, S. Busche, M. Caron, S. H. Chen, W. Cheung, O. Delaneau, E. T. Dermitzakis, H. Elding, I. Colgiu, F. O. Bagger, P. Flicek, E. Habibi, V. Iotchkova, E. Janssen-Megens, B. Kim, H. Lehrach, E. Lowy, A. Mandoli, F. Matarese, M. T. Maurano, J. A. Morris, V. Pancaldi, F. Pourfarzad, K. Rehnstrom, A. Rendon, T. Risch, N. Sharifi, M. M. Simon, M. Sultan, A. Valencia, K. Walter, S. Y. Wang, M. Frontini, S. E. Antonarakis, L. Clarke, M. L. Yaspo, S. Beck, R. Guigo, D. Rico, J. H. A. Martens, W. H. Ouwehand, T. W. Kuijpers, D. S. Paul, H. G. Stunnenberg, O. Stegle, K. Downes, T. Pastinen and N. Soranzo (2016). “Genetic Drivers of Epigenetic and Transcriptional Variation in Human Immune Cells.” Cell 167(5): 1398–1414.e1324.

Chiou, J., R. J. Geusz, M.-L. Okino, J. Y. Han, M. Miller, R. Melton, E. Beebe, P. Benaglio, S. Huang, K. Korgaonkar, S. Heller, A. Kleger, S. Preissl, D. U. Gorkin, M. Sander and K. J. Gaulton (2021). “Interpreting type 1 diabetes risk with genetics and single-cell epigenomics.” Nature 594(7863): 398– 402.

Chun, S., A. Casparino, N. A. Patsopoulos, D. C. Croteau-Chonka, B. A. Raby, P. L. De Jager, S. R. Sunyaev and C. Cotsapas (2017). “Limited statistical evidence for shared genetic effects of eQTLs and autoimmune-disease-associated loci in three major immune-cell types.” Nat Genet 49(4): 600–605.

Connally, N. J., S. Nazeen, D. Lee, H. Shi, J. Stamatoyannopoulos, S. Chun, C. Cotsapas, C. A. Cassa and S. R. Sunyaev (2022). “The missing link between genetic association and regulatory function.” Elife 11.

Datlinger, P., A. F. Rendeiro, C. Schmidl, T. Krausgruber, P. Traxler, J. Klughammer, L. C. Schuster, A. Kuchler, D. Alpar and C. Bock (2017). “Pooled CRISPR screening with single-cell transcriptome readout.” Nat Methods 14(3): 297–301.

Degner, J. F., A. A. Pai, R. Pique-Regi, J. B. Veyrieras, D. J. Gaffney, J. K. Pickrell, S. De Leon, K. Michelini, N. Lewellen, G. E. Crawford, M. Stephens, Y. Gilad and J. K. Pritchard (2012). “DNase I sensitivity QTLs are a major determinant of human expression variation.” Nature 482(7385): 390–394.

Diao, Y., B. Li, Z. Meng, I. Jung, A. Y. Lee, J. Dixon, L. Maliskova, K. L. Guan, Y. Shen and B. Ren (2016). “A new class of temporarily phenotypic enhancers identified by CRISPR/Cas9-mediated genetic screening.” Genome Res 26(3): 397–405.

Donovan, M. K. R., A. D’Antonio-Chronowska, M. D’Antonio and K. A. Frazer (2020). “Cellular deconvolution of GTEx tissues powers discovery of disease and cell-type associated regulatory variants.” Nat Commun 11(1): 955.

Durinck, S., Y. Moreau, A. Kasprzyk, S. Davis, B. De Moor, A. Brazma and W. Huber (2005). “BioMart and Bioconductor: a powerful link between biological databases and microarray data analysis.” Bioinformatics 21(16): 3439–3440.

Durinck, S., P.sss T. Spellman, E. Birney and W. Huber (2009). “Mapping identifiers for the integration of genomic datasets with the R/Bioconductor package biomaRt.” Nat Protoc 4(8): 1184–1191.

ENCODE-Project-Consortium (2012). “An integrated encyclopedia of DNA elements in the human genome.” Nature 489(7414): 57–74.

Farh, K. K., A. Marson, J. Zhu, M. Kleinewietfeld, W. J. Housley, S. Beik, N. Shoresh, H. Whitton, R. J. Ryan, A. A. Shishkin, M. Hatan, M. J. Carrasco-Alfonso, D. Mayer, C. J. Luckey, N. A. Patsopoulos, P. L. De Jager, V. K. Kuchroo, C. B. Epstein, M. J. Daly, D. A. Hafler and B. E. Bernstein (2015). “Genetic and epigenetic fine mapping of causal autoimmune disease variants.” Nature 518(7539): 337–343.

Festenstein, R., M. Tolaini, P. Corbella, C. Mamalaki, J. Parrington, M. Fox, A. Miliou, M. Jones and D. Kioussis (1996). “Locus control region function and heterochromatin-induced position effect variegation.” Science 271(5252): 1123–1125.

Finak, G., A. McDavid, M. Yajima, J. Deng, V. Gersuk, A. K. Shalek, C. K. Slichter, H. W. Miller, M. J. McElrath, M. Prlic, P. S. Linsley and R. Gottardo (2015). “MAST: a flexible statistical framework for assessing transcriptional changes and characterizing heterogeneity in single-cell RNA sequencing data.” Genome Biol 16: 278.

Finan, C., A. Gaulton, F. A. Kruger, R. T. Lumbers, T. Shah, J. Engmann, L. Galver, R. Kelley, A. Karlsson, R. Santos, J. P. Overington, A. D. Hingorani and J. P. Casas (2017). “The druggable genome and support for target identification and validation in drug development.” Sci Transl Med 9(383).

Finucane, H. K., Y. A. Reshef, V. Anttila, K. Slowikowski, A. Gusev, A. Byrnes, S. Gazal, P. R. Loh, C. Lareau, N. Shoresh, G. Genovese, A. Saunders, E. Macosko, S. Pollack, J. R. B. Perry, J. D. Buenrostro, B. E. Bernstein, S. Raychaudhuri, S. McCarroll, B. M. Neale and A. L. Price (2018). “Heritability enrichment of specifically expressed genes identifies disease-relevant tissues and cell types.” Nat Genet 50(4): 621–629.

Fletcher, M. (2023). “Linking GWAS to gene regulation.” Nat Genet 55(2): 167.

Freimer, J. W., O. Shaked, S. Naqvi, N. Sinnott-Armstrong, A. Kathiria, A. F. Chen, J. T. Cortez, W. J. Greenleaf, J. K. Pritchard and A. Marson (2021). “Systematic discovery and perturbation of regulatory genes in human T cells reveals the architecture of immune networks.” bioRxiv: 2021.2004.2018.440363.

French, J. D. and S. L. Edwards (2020). “The Role of Noncoding Variants in Heritable Disease.” Trends Genet 36(11): 880–891.

Fulco, C. P., M. Munschauer, R. Anyoha, G. Munson, S. R. Grossman, E. M. Perez, M. Kane, B. Cleary, E. S. Lander and J. M. Engreitz (2016). “Systematic mapping of functional enhancer-promoter connections with CRISPR interference.” Science 354(6313): 769–773.

Fulco, C. P., J. Nasser, T. R. Jones, G. Munson, D. T. Bergman, V. Subramanian, S. R. Grossman, R. Anyoha, B. R. Doughty, T. A. Patwardhan, T. H. Nguyen, M. Kane, E. M. Perez, N. C. Durand, C. A. Lareau, E. K. Stamenova, E. L. Aiden, E. S. Lander and J. M. Engreitz (2019). “Activity-by-contact model of enhancer-promoter regulation from thousands of CRISPR perturbations.” Nature genetics 51(12): 1664–1669.

Gasperini, M., G. M. Findlay, A. McKenna, J. H. Milbank, C. Lee, M. D. Zhang, D. A. Cusanovich and J. Shendure (2017). “CRISPR/Cas9-Mediated Scanning for Regulatory Elements Required for HPRT1 Expression via Thousands of Large, Programmed Genomic Deletions.” Am J Hum Genet 101(2): 192–205.

Gasperini, M., A. J. Hill, J. L. McFaline-Figueroa, B. Martin, S. Kim, M. D. Zhang, D. Jackson, A. Leith, J. Schreiber, W. S. Noble, C. Trapnell, N. Ahituv and J. Shendure (2019). “A Genome-wide Framework for Mapping Gene Regulation via Cellular Genetic Screens.” Cell 176(1-2): 377–390.e319.

Gasperini, M., J. M. Tome and J. Shendure (2020). “Towards a comprehensive catalogue of validated and target-linked human enhancers.” Nature reviews. Genetics 21(5): 292–310.

Gate, R. E., M. C. Kim, A. Lu, D. Lee, E. Shifrut, M. Subramaniam, A. Marson and C. J. Ye (2019). “Mapping gene regulatory networks of primary CD4+ T cells using single-cell genomics and genome engineering.” bioRxiv: 678060.

Gaulton, K. J., T. Ferreira, Y. Lee, A. Raimondo, R. Mägi, M. E. Reschen, A. Mahajan, A. Locke, N. W. Rayner, N. Robertson, R. A. Scott, I. Prokopenko, L. J. Scott, T. Green, T. Sparso, D. Thuillier, L. Yengo, H. Grallert, S. Wahl, M. Frånberg, R. J. Strawbridge, H. Kestler, H. Chheda, L. Eisele, S. Gustafsson, V. Steinthorsdottir, G. Thorleifsson, L. Qi, L. C. Karssen, E. M. van Leeuwen, S. M. Willems, M. Li, H. Chen, C. Fuchsberger, P. Kwan, C. Ma, M. Linderman, Y. Lu, S. K. Thomsen, J. K. Rundle, N. L. Beer, M. van de Bunt, A. Chalisey, H. M. Kang, B. F. Voight, G. R. Abecasis, P. Almgren, D. Baldassarre, B. Balkau, R. Benediktsson, M. Blüher, H. Boeing, L. L. Bonnycastle, E. P. Bottinger, N. P. Burtt, J. Carey, G. Charpentier, P. S. Chines, M. C. Cornelis, D. J. Couper, A. T. Crenshaw, R. M. van Dam, A. S. Doney, M. Dorkhan, S. Edkins, J. G. Eriksson, T. Esko, E. Eury, J. Fadista, J. Flannick, P. Fontanillas, C. Fox, P. W. Franks, K. Gertow, C. Gieger, B. Gigante, O. Gottesman, G. B. Grant, N. Grarup, C. J. Groves, M. Hassinen, C. T. Have, C. Herder, O. L. Holmen, A. B. Hreidarsson, S. E. Humphries, D. J. Hunter, A. U. Jackson, A. Jonsson, M. E. Jørgensen, T. Jørgensen, W. H. Kao, N. D. Kerrison, L. Kinnunen, N. Klopp, A. Kong, P. Kovacs, P. Kraft, J. Kravic, C. Langford, K. Leander, L. Liang, P. Lichtner, C. M. Lindgren, E. Lindholm, A. Linneberg, C. T. Liu, S. Lobbens, J. Luan, V. Lyssenko, S. Männistö, O. McLeod, J. Meyer, E. Mihailov, G. Mirza, T. W. Mühleisen, M. Müller-Nurasyid, C. Navarro, M. M. Nöthen, N. N. Oskolkov, K. R. Owen, D. Palli, S. Pechlivanis, L. Peltonen, J. R. Perry, C. G. Platou, M. Roden, D. Ruderfer, D. Rybin, Y. T. van der Schouw, B. Sennblad, G. Sigurðsson, A. Stančáková, G. Steinbach, P. Storm, K. Strauch, H. M. Stringham, Q. Sun, B. Thorand, E. Tikkanen, A. Tonjes, J. Trakalo, E. Tremoli, T. Tuomi, R. Wennauer, S. Wiltshire, A. R. Wood, E. Zeggini, I. Dunham, E. Birney, L. Pasquali, J. Ferrer, R. J. Loos, J. Dupuis, J. C. Florez, E. Boerwinkle, J. S. Pankow, C. van Duijn, E. Sijbrands, J. B. Meigs, F. B. Hu, U. Thorsteinsdottir, K. Stefansson, T. A. Lakka, R. Rauramaa, M. Stumvoll, N. L. Pedersen, L. Lind, S. M. Keinanen-Kiukaanniemi, E. Korpi-Hyövälti, T. E. Saaristo, J. Saltevo, J. Kuusisto, M. Laakso, A. Metspalu, R. Erbel, K. H. Jöcke, S. Moebus, S. Ripatti, V. Salomaa, E. Ingelsson, B. O. Boehm, R. N. Bergman, F. S. Collins, K. L. Mohlke, H. Koistinen, J. Tuomilehto, K. Hveem, I. Njølstad, P. Deloukas, P. J. Donnelly, T. M. Frayling, A. T. Hattersley, U. de Faire, A. Hamsten, T. Illig, A. Peters, S. Cauchi, R. Sladek, P. Froguel, T. Hansen, O. Pedersen, A. D. Morris, C. N. Palmer, S. Kathiresan, O. Melander, P. M. Nilsson, L. C. Groop, I. Barroso, C. Langenberg, N. J. Wareham, C. A. O’Callaghan, A. L. Gloyn, D. Altshuler, M. Boehnke, T. M. Teslovich, M. I. McCarthy and A. P. Morris (2015). “Genetic fine mapping and genomic annotation defines causal mechanisms at type 2 diabetes susceptibility loci.” Nat Genet 47(12): 1415–1425.

Germain, P. L., A. Lun, C. Garcia Meixide, W. Macnair and M. D. Robinson (2021). “Doublet identification in single-cell sequencing data using scDblFinder.” F1000Res 10: 979.

Giambartolomei, C., D. Vukcevic, E. E. Schadt, L. Franke, A. D. Hingorani, C. Wallace and V. Plagnol (2014). “Bayesian test for colocalisation between pairs of genetic association studies using summary statistics.” PLoS Genet 10(5): e1004383.

Gilbert, L. A., M. A. Horlbeck, B. Adamson, J. E. Villalta, Y. Chen, E. H. Whitehead, C. Guimaraes, B. Panning, H. L. Ploegh, M. C. Bassik, L. S. Qi, M. Kampmann and J. S. Weissman (2014). “Genome-Scale CRISPR-Mediated Control of Gene Repression and Activation.” Cell 159(3): 647–661.

Griffiths, J. A., A. C. Richard, K. Bach, A. T. L. Lun and J. C. Marioni (2018). “Detection and removal of barcode swapping in single-cell RNA-seq data.” Nat Commun 9(1): 2667.

GTEX-Consortium (2020). “The GTEx Consortium atlas of genetic regulatory effects across human tissues.” Science 369(6509): 1318–1330.

Hill, A. J., J. L. McFaline-Figueroa, L. M. Starita, M. J. Gasperini, K. A. Matreyek, J. Packer, D. Jackson, J. Shendure and C. Trapnell (2018). “On the design of CRISPR-based single-cell molecular screens.” Nat Methods 15(4): 271–274.

Javierre, B. M., O. S. Burren, S. P. Wilder, R. Kreuzhuber, S. M. Hill, S. Sewitz, J. Cairns, S. W. Wingett, C. Várnai, M. J. Thiecke, F. Burden, S. Farrow, A. J. Cutler, K. Rehnström, K. Downes, L. Grassi, M. Kostadima, P. Freire-Pritchett, F. Wang, H. G. Stunnenberg, J. A. Todd, D. R. Zerbino, O. Stegle, W. H. Ouwehand, M. Frontini, C. Wallace, M. Spivakov and P. Fraser (2016). “Lineage-Specific Genome Architecture Links Enhancers and Non-coding Disease Variants to Target Gene Promoters.” Cell 167(5): 1369–1384.e1319.

Joung, J., S. Konermann, J. S. Gootenberg, O. O. Abudayyeh, R. J. Platt, M. D. Brigham, N. E. Sanjana and F. Zhang (2017). “Genome-scale CRISPR-Cas9 knockout and transcriptional activation screening.” Nat Protoc 12(4): 828–863.

Kaptein, L. C., M. Breuer, D. Valerio and V. W. van Beusechem (1998). “Expression pattern of CD2 locus control region containing retroviral vectors in hemopoietic cells in vitro and in vivo.” Gene Ther 5(3): 320–330.

Kaur, G. and J. M. Dufour (2012). “Cell lines: Valuable tools or useless artifacts.” Spermatogenesis 2(1): 1–5.

Kim-Hellmuth, S., F. Aguet, M. Oliva, M. Muñoz-Aguirre, S. Kasela, V. Wucher, S. E. Castel, A. R. Hamel, A. Viñuela, A. L. Roberts, S. Mangul, X. Wen, G. Wang, A. N. Barbeira, D. Garrido-Martín, B. B. Nadel, Y. Zou, R. Bonazzola, J. Quan, A. Brown, A. Martinez-Perez, J. M. Soria, G. Getz, E. T. Dermitzakis, K. S. Small, M. Stephens, H. S. Xi, H. K. Im, R. Guigó, A. V. Segrè, B. E. Stranger, K. G. Ardlie and T. Lappalainen (2020). “Cell type-specific genetic regulation of gene expression across human tissues.” Science 369(6509).

King, E. A., J. W. Davis and J. F. Degner (2019). “Are drug targets with genetic support twice as likely to be approved? Revised estimates of the impact of genetic support for drug mechanisms on the probability of drug approval.” PLoS Genet 15(12): e1008489.

Klann, T. S., J. B. Black, M. Chellappan, A. Safi, L. Song, I. B. Hilton, G. E. Crawford, T. E. Reddy and C. A. Gersbach (2017). “CRISPR-Cas9 epigenome editing enables high-throughput screening for functional regulatory elements in the human genome.” Nature biotechnology 35(6): 561–568.

Korkmaz, G., R. Lopes, A. P. Ugalde, E. Nevedomskaya, R. Han, K. Myacheva, W. Zwart, R. Elkon and R. Agami (2016). “Functional genetic screens for enhancer elements in the human genome using CRISPR-Cas9.” Nat Biotechnol 34(2): 192–198.

Kundaje, A., W. Meuleman, J. Ernst, M. Bilenky, A. Yen, A. Heravi-Moussavi, P. Kheradpour, Z. Zhang, J. Wang, M. J. Ziller, V. Amin, J. W. Whitaker, M. D. Schultz, L. D. Ward, A. Sarkar, G. Quon, R. S. Sandstrom, M. L. Eaton, Y. C. Wu, A. R. Pfenning, X. Wang, M. Claussnitzer, Y. Liu, C. Coarfa, R. A. Harris, N. Shoresh, C. B. Epstein, E. Gjoneska, D. Leung, W. Xie, R. D. Hawkins, R. Lister, C. Hong, P. Gascard, A. J. Mungall, R. Moore, E. Chuah, A. Tam, T. K. Canfield, R. S. Hansen, R. Kaul, P. J. Sabo, M. S. Bansal, A. Carles, J. R. Dixon, K. H. Farh, S. Feizi, R. Karlic, A. R. Kim, A. Kulkarni, D. Li, R. Lowdon, G. Elliott, T. R. Mercer, S. J. Neph, V. Onuchic, P. Polak, N. Rajagopal, P. Ray, R. C. Sallari, K. T. Siebenthall, N. A. Sinnott-Armstrong, M. Stevens, R. E. Thurman, J. Wu, B. Zhang, X. Zhou, A. E. Beaudet, L. A. Boyer, P. L. De Jager, P. J. Farnham, S. J. Fisher, D. Haussler, S. J. Jones, W. Li, M. A. Marra, M. T. McManus, S. Sunyaev, J. A. Thomson, T. D. Tlsty, L. H. Tsai, W. Wang, R. A. Waterland, M. Q. Zhang, L. H. Chadwick, B. E. Bernstein, J. F. Costello, J. R. Ecker, M. Hirst, A. Meissner, A. Milosavljevic, B. Ren, J. A. Stamatoyannopoulos, T. Wang and M. Kellis (2015). “Integrative analysis of 111 reference human epigenomes.” Nature 518(7539): 317–330.

Lake, R. A., D. Wotton and M. J. Owen (1990). “A 3’ transcriptional enhancer regulates tissue-specific expression of the human CD2 gene.” Embo j 9(10): 3129–3136.

Law, C. W., Y. Chen, W. Shi and G. K. Smyth (2014). “voom: Precision weights unlock linear model analysis tools for RNA-seq read counts.” Genome Biol 15(2): R29.

Li, K., Y. Liu, H. Cao, Y. Zhang, Z. Gu, X. Liu, A. Yu, P. Kaphle, K. E. Dickerson, M. Ni and J. Xu (2020). “Interrogation of enhancer function by enhancer-targeting CRISPR epigenetic editing.” Nat Commun 11(1): 485.

Lun, A. (2022). “metapod: Meta-Analyses on P-Values of Differential Analyses. R package version 1.6.0.”, from https://www.bioconductor.org/packages/release/bioc/html/metapod.html.

Lun, A. T., D. J. McCarthy and J. C. Marioni (2016). “A step-by-step workflow for low-level analysis of single-cell RNA-seq data with Bioconductor.” F1000Res 5: 2122.

Lun, A. T. L., S. Riesenfeld, T. Andrews, T. P. Dao, T. Gomes and J. C. Marioni (2019). “EmptyDrops: distinguishing cells from empty droplets in droplet-based single-cell RNA sequencing data.” Genome Biol 20(1): 63.

Maller, J. B., G. McVean, J. Byrnes, D. Vukcevic, K. Palin, Z. Su, J. M. Howson, A. Auton, S. Myers, A. Morris, M. Pirinen, M. A. Brown, P. R. Burton, M. J. Caulfield, A. Compston, M. Farrall, A. S. Hall, A. T. Hattersley, A. V. Hill, C. G. Mathew, M. Pembrey, J. Satsangi, M. R. Stratton, J. Worthington, N. Craddock, M. Hurles, W. Ouwehand, M. Parkes, N. Rahman, A. Duncanson, J. A. Todd, D. P. Kwiatkowski, N. J. Samani, S. C. Gough, M. I. McCarthy, P. Deloukas and P. Donnelly (2012). “Bayesian refinement of association signals for 14 loci in 3 common diseases.” Nat Genet 44(12): 1294–1301.

Maurano, M. T., R. Humbert, E. Rynes, R. E. Thurman, E. Haugen, H. Wang, A. P. Reynolds, R. Sandstrom, H. Qu, J. Brody, A. Shafer, F. Neri, K. Lee, T. Kutyavin, S. Stehling-Sun, A. K. Johnson, T. K. Canfield, E. Giste, M. Diegel, D. Bates, R. S. Hansen, S. Neph, P. J. Sabo, S. Heimfeld, A. Raubitschek, S. Ziegler, C. Cotsapas, N. Sotoodehnia, I. Glass, S. R. Sunyaev, R. Kaul and J. A. Stamatoyannopoulos (2012). “Systematic localization of common disease-associated variation in regulatory DNA.” Science 337(6099): 1190–1195.

McCarthy, D. J., K. R. Campbell, A. T. Lun and Q. F. Wills (2017). “Scater: pre-processing, quality control, normalization and visualization of single-cell RNA-seq data in R.” Bioinformatics 33(8): 1179–1186.

Meuleman, W., A. Muratov, E. Rynes, J. Halow, K. Lee, D. Bates, M. Diegel, D. Dunn, F. Neri, A. Teodosiadis, A. Reynolds, E. Haugen, J. Nelson, A. Johnson, M. Frerker, M. Buckley, R. Sandstrom, J. Vierstra, R. Kaul and J. Stamatoyannopoulos (2020). “Index and biological spectrum of human DNase I hypersensitive sites.” Nature 584(7820): 244–251.

Montefiori, L. E., D. R. Sobreira, N. J. Sakabe, I. Aneas, A. C. Joslin, G. T. Hansen, G. Bozek, I. P. Moskowitz, E. M. McNally and M. A. Nóbrega (2018). “A promoter interaction map for cardiovascular disease genetics.” eLife 7.

Moore, J. E., M. J. Purcaro, H. E. Pratt, C. B. Epstein, N. Shoresh, J. Adrian, T. Kawli, C. A. Davis, A. Dobin, R. Kaul, J. Halow, E. L. Van Nostrand, P. Freese, D. U. Gorkin, Y. Shen, Y. He, M. Mackiewicz, F. Pauli-Behn, B. A. Williams, A. Mortazavi, C. A. Keller, X. O. Zhang, S. I. Elhajjajy, J. Huey, D. E. Dickel, V. Snetkova, X. Wei, X. Wang, J. C. Rivera-Mulia, J. Rozowsky, J. Zhang, S. B. Chhetri, J. Zhang, A. Victorsen, K. P. White, A. Visel, G. W. Yeo, C. B. Burge, E. Lécuyer, D. M. Gilbert, J. Dekker, J. Rinn, E. M. Mendenhall, J. R. Ecker, M. Kellis, R. J. Klein, W. S. Noble, A. Kundaje, R. Guigó, P. J. Farnham, J. M. Cherry, R. M. Myers, B. Ren, B. R. Graveley, M. B. Gerstein, L. A. Pennacchio, M. P. Snyder, B. E. Bernstein, B. Wold, R. C. Hardison, T. R. Gingeras, J. A. Stamatoyannopoulos and Z. Weng (2020). “Expanded encyclopaedias of DNA elements in the human and mouse genomes.” Nature 583(7818): 699–710.

Morris, J. A., C. Caragine, Z. Daniloski, J. Domingo, T. Barry, L. Lu, K. Davis, M. Ziosi, D. A. Glinos, S. Hao, E. P. Mimitou, P. Smibert, K. Roeder, E. Katsevich, T. Lappalainen and N. E. Sanjana (2023). “Discovery of target genes and pathways at GWAS loci by pooled single-cell CRISPR screens.” Science: eadh7699.

Mostafavi, H., J. P. Spence, S. Naqvi and J. K. Pritchard (2022). “Limited overlap of eQTLs and GWAS hits due to systematic differences in discovery.” bioRxiv: 2022.2005.2007.491045.

Mumbach, M. R., A. T. Satpathy, E. A. Boyle, C. Dai, B. G. Gowen, S. W. Cho, M. L. Nguyen, A. J. Rubin, J. M. Granja, K. R. Kazane, Y. Wei, T. Nguyen, P. G. Greenside, M. R. Corces, J. Tycko, D. R. Simeonov, N. Suliman, R. Li, J. Xu, R. A. Flynn, A. Kundaje, P. A. Khavari, A. Marson, J. E. Corn, T. Quertermous, W. J. Greenleaf and H. Y. Chang (2017). “Enhancer connectome in primary human cells identifies target genes of disease-associated DNA elements.” Nat Genet 49(11): 1602–1612.

Nasser, J., D. T. Bergman, C. P. Fulco, P. Guckelberger, B. R. Doughty, T. A. Patwardhan, T. R. Jones, T. H. Nguyen, J. C. Ulirsch, F. Lekschas, K. Mualim, H. M. Natri, E. M. Weeks, G. Munson, M. Kane, H. Y. Kang, A. Cui, J. P. Ray, T. M. Eisenhaure, R. L. Collins, K. Dey, H. Pfister, A. L. Price, C. B. Epstein, A. Kundaje, R. J. Xavier, M. J. Daly, H. Huang, H. K. Finucane, N. Hacohen, E. S. Lander and J. M. Engreitz (2021). “Genome-wide enhancer maps link risk variants to disease genes.” Nature 593(7858): 238–243.

Nathan, A., S. Asgari, K. Ishigaki, T. Amariuta, Y. Luo, J. I. Beynor, Y. Baglaenko, S. Suliman, A. Price, L. Lecca, M. B. Murray, D. B. Moody and S. Raychaudhuri (2021). “Modeling memory T cell states at single-cell resolution identifies <em>in vivo</em> state-dependence of eQTLs influencing disease.” bioRxiv: 2021.2007.2029.454316.

Nathan, A., S. Asgari, K. Ishigaki, C. Valencia, T. Amariuta, Y. Luo, J. I. Beynor, Y. Baglaenko, S. Suliman, A. L. Price, L. Lecca, M. B. Murray, D. B. Moody and S. Raychaudhuri (2022). “Single-cell eQTL models reveal dynamic T cell state dependence of disease loci.” Nature 606(7912): 120–128.

Nelson, M. R., H. Tipney, J. L. Painter, J. Shen, P. Nicoletti, Y. Shen, A. Floratos, P. C. Sham, M. J. Li, J. Wang, L. R. Cardon, J. C. Whittaker and P. Sanseau (2015). “The support of human genetic evidence for approved drug indications.” Nat Genet 47(8): 856–860.

Nott, A., I. R. Holtman, N. G. Coufal, J. C. M. Schlachetzki, M. Yu, R. Hu, C. Z. Han, M. Pena, J. Xiao, Y. Wu, Z. Keulen, M. P. Pasillas, C. O’Connor, C. K. Nickl, S. T. Schafer, Z. Shen, R. A. Rissman, J. B. Brewer, D. Gosselin, D. D. Gonda, M. L. Levy, M. G. Rosenfeld, G. McVicker, F. H. Gage, B. Ren and C. K. Glass (2019). “Brain cell type-specific enhancer-promoter interactome maps and disease-risk association.” Science (New York, N.Y.) 366(6469): 1134–1139.

Ochoa, D., M. Karim, M. Ghoussaini, D. G. Hulcoop, E. M. McDonagh and I. Dunham (2022). “Human genetics evidence supports two-thirds of the 2021 FDA-approved drugs.” Nat Rev Drug Discov 21(8): 551.

Okada, Y., D. Wu, G. Trynka, T. Raj, C. Terao, K. Ikari, Y. Kochi, K. Ohmura, A. Suzuki, S. Yoshida, R. R. Graham, A. Manoharan, W. Ortmann, T. Bhangale, J. C. Denny, R. J. Carroll, A. E. Eyler, J. D. Greenberg, J. M. Kremer, D. A. Pappas, L. Jiang, J. Yin, L. Ye, D. F. Su, J. Yang, G. Xie, E. Keystone, H. J. Westra, T. Esko, A. Metspalu, X. Zhou, N. Gupta, D. Mirel, E. A. Stahl, D. Diogo, J. Cui, K. Liao, M. H. Guo, K. Myouzen, T. Kawaguchi, M. J. Coenen, P. L. van Riel, M. A. van de Laar, H. J. Guchelaar, T. W. Huizinga, P. Dieudé, X. Mariette, S. L. Bridges, Jr., A. Zhernakova, R. E. Toes, P. P. Tak, C. Miceli-Richard, S. Y. Bang, H. S. Lee, J. Martin, M. A. Gonzalez-Gay, L. Rodriguez-Rodriguez, S. Rantapää-Dahlqvist, L. Arlestig, H. K. Choi, Y. Kamatani, P. Galan, M. Lathrop, S. Eyre, J. Bowes, A. Barton, N. de Vries, L. W. Moreland, L. A. Criswell, E. W. Karlson, A. Taniguchi, R. Yamada, M. Kubo, J. S. Liu, S. C. Bae, J. Worthington, L. Padyukov, L. Klareskog, P. K. Gregersen, S. Raychaudhuri, B. E. Stranger, P. L. De Jager, L. Franke, P. M. Visscher, M. A. Brown, H. Yamanaka, T. Mimori, A. Takahashi, H. Xu, T. W. Behrens, K. A. Siminovitch, S. Momohara, F. Matsuda, K. Yamamoto and R. M. Plenge (2014). “Genetics of rheumatoid arthritis contributes to biology and drug discovery.” Nature 506(7488): 376–381.

Ota, M., Y. Nagafuchi, H. Hatano, K. Ishigaki, C. Terao, Y. Takeshima, H. Yanaoka, S. Kobayashi, M. Okubo, H. Shirai, Y. Sugimori, J. Maeda, M. Nakano, S. Yamada, R. Yoshida, H. Tsuchiya, Y. Tsuchida, S. Akizuki, H. Yoshifuji, K. Ohmura, T. Mimori, K. Yoshida, D. Kurosaka, M. Okada, K. Setoguchi, H. Kaneko, N. Ban, N. Yabuki, K. Matsuki, H. Mutoh, S. Oyama, M. Okazaki, H. Tsunoda, Y. Iwasaki, S. Sumitomo, H. Shoda, Y. Kochi, Y. Okada, K. Yamamoto, T. Okamura and K. Fujio (2021). “Dynamic landscape of immune cell-specific gene regulation in immune-mediated diseases.” Cell 184(11): 3006–3021.e3017.

Pastor, D. M., L. S. Poritz, T. L. Olson, C. L. Kline, L. R. Harris, W. A. Koltun, V. M. Chinchilli and R. B. Irby (2010). “Primary cell lines: false representation or model system? a comparison of four human colorectal tumors and their coordinately established cell lines.” Int J Clin Exp Med 3(1): 69–83.

Paternoster, L., M. Standl, J. Waage, H. Baurecht, M. Hotze, D. P. Strachan, J. A. Curtin, K. Bønnelykke, C. Tian, A. Takahashi, J. Esparza-Gordillo, A. C. Alves, J. P. Thyssen, H. T. den Dekker, M. A. Ferreira, E. Altmaier, P. M. Sleiman, F. L. Xiao, J. R. Gonzalez, I. Marenholz, B. Kalb, M. P. Yanes, C. J. Xu, L. Carstensen, M. M. Groen-Blokhuis, C. Venturini, C. E. Pennell, S. J. Barton, A. M. Levin, I. Curjuric, M. Bustamante, E. Kreiner-Møller, G. A. Lockett, J. Bacelis, S. Bunyavanich, R. A. Myers, A. Matanovic, A. Kumar, J. Y. Tung, T. Hirota, M. Kubo, W. L. McArdle, A. J. Henderson, J. P. Kemp, J. Zheng, G. D. Smith, F. Rüschendorf, A. Bauerfeind, M. A. Lee-Kirsch, A. Arnold, G. Homuth, C. O. Schmidt, E. Mangold, S. Cichon, T. Keil, E. Rodríguez, A. Peters, A. Franke, W. Lieb, N. Novak, R. Fölster-Holst, M. Horikoshi, J. Pekkanen, S. Sebert, L. L. Husemoen, N. Grarup, J. C. de Jongste, F. Rivadeneira, A. Hofman, V. W. Jaddoe, S. G. Pasmans, N. J. Elbert, A. G. Uitterlinden, G. B. Marks, P. J. Thompson, M. C. Matheson, C. F. Robertson, J. S. Ried, J. Li, X. B. Zuo, X. D. Zheng, X. Y. Yin, L. D. Sun, M. A. McAleer, G. M. O’Regan, C. M. Fahy, L. E. Campbell, M. Macek, M. Kurek, D. Hu, C. Eng, D. S. Postma, B. Feenstra, F. Geller, J. J. Hottenga, C. M. Middeldorp, P. Hysi, V. Bataille, T. Spector, C. M. Tiesler, E. Thiering, B. Pahukasahasram, J. J. Yang, M. Imboden, S. Huntsman, N. Vilor-Tejedor, C. L. Relton, R. Myhre, W. Nystad, A. Custovic, S. T. Weiss, D. A. Meyers, C. Söderhäll, E. Melén, C. Ober, B. A. Raby, A. Simpson, B. Jacobsson, J. W. Holloway, H. Bisgaard, J. Sunyer, N. M. P. Hensch, L. K. Williams, K. M. Godfrey, C. A. Wang, D. I. Boomsma, M. Melbye, G. H. Koppelman, D. Jarvis, W. I. McLean, A. D. Irvine, X. J. Zhang, H. Hakonarson, C. Gieger, E. G. Burchard, N. G. Martin, L. Duijts, A. Linneberg, M. R. Jarvelin, M. M. Noethen, S. Lau, N. Hübner, Y. A. Lee, M. Tamari, D. A. Hinds, D. Glass, S. J. Brown, J. Heinrich, D. M. Evans and S. Weidinger (2015). “Multi-ancestry genome-wide association study of 21,000 cases and 95,000 controls identifies new risk loci for atopic dermatitis.” Nat Genet 47(12): 1449–1456.

Patsopoulos, N. A., S. E. Baranzini, A. Santaniello, P. Shoostari, C. Cotsapas, G. Wong, A. H. Beecham, T. James, J. Replogle, I. S. Vlachos, C. McCabe, T. H. Pers, A. Brandes, C. White, B. Keenan, M. Cimpean, P. Winn, I.-P. Panteliadis, A. Robbins, T. F. M. Andlauer, O. Zarzycki, B. Dubois, A. Goris, H. B. Søndergaard, F. Sellebjerg, P. S. Sorensen, H. Ullum, L. W. Thørner, J. Saarela, I. Cournu-Rebeix, V. Damotte, B. Fontaine, L. Guillot-Noel, M. Lathrop, S. Vukusic, A. Berthele, V. Pongratz, D. Buck, C. Gasperi, C. Graetz, V. Grummel, B. Hemmer, M. Hoshi, B. Knier, T. Korn, C. M. Lill, F. Luessi, M. Mühlau, F. Zipp, E. Dardiotis, C. Agliardi, A. Amoroso, N. Barizzone, M. D. Benedetti, L. Bernardinelli, P. Cavalla, F. Clarelli, G. Comi, D. Cusi, F. Esposito, L. Ferrè, D. Galimberti, C. Guaschino, M. A. Leone, V. Martinelli, L. Moiola, M. Salvetti, M. Sorosina, D. Vecchio, A. Zauli, S. Santoro, N. Mancini, M. Zuccalà, J. Mescheriakova, C. van Duijn, S. D. Bos, E. G. Celius, A. Spurkland, M. Comabella, X. Montalban, L. Alfredsson, I. L. Bomfim, D. Gomez-Cabrero, J. Hillert, M. Jagodic, M. Lindén, F. Piehl, I. Jelčić, R. Martin, M. Sospedra, A. Baker, M. Ban, C. Hawkins, P. Hysi, S. Kalra, F. Karpe, J. Khadake, G. Lachance, P. Molyneux, M. Neville, J. Thorpe, E. Bradshaw, S. J. Caillier, P. Calabresi, B. A. C. Cree, A. Cross, M. Davis, P. W. I. de Bakker, S. Delgado, M. Dembele, K. Edwards, K. Fitzgerald, I. Y. Frohlich, P.-A. Gourraud, J. L. Haines, H. Hakonarson, D. Kimbrough, N. Isobe, I. Konidari, E. Lathi, M. H. Lee, T. Li, D. An, A. Zimmer, L. Madireddy, C. P. Manrique, M. Mitrovic, M. Olah, E. Patrick, M. A. Pericak-Vance, L. Piccio, C. Schaefer, H. Weiner, K. Lage, A. Compston, D. Hafler, H. F. Harbo, S. L. Hauser, G. Stewart, S. D’Alfonso, G. Hadjigeorgiou, B. Taylor, L. F. Barcellos, D. Booth, R. Hintzen, I. Kockum, F. Martinelli-Boneschi, J. L. McCauley, J. R. Oksenberg, A. Oturai, S. Sawcer, A. J. Ivinson, T. Olsson and P. L. De Jager (2019). “Multiple sclerosis genomic map implicates peripheral immune cells and microglia in susceptibility.” Science 365(6460): eaav7188.

Pritchard, J. E., T. A. O’Mara and D. M. Glubb (2017). “Enhancing the Promise of Drug Repositioning through Genetics.” Front Pharmacol 8: 896.

Rajagopal, N., S. Srinivasan, K. Kooshesh, Y. Guo, M. D. Edwards, B. Banerjee, T. Syed, B. J. Emons, D. K. Gifford and R. I. Sherwood (2016). “High-throughput mapping of regulatory DNA.” Nat Biotechnol 34(2): 167–174.

Sakaguchi, S., N. Mikami, J. B. Wing, A. Tanaka, K. Ichiyama and N. Ohkura (2020). “Regulatory T Cells and Human Disease.” Annu Rev Immunol 38: 541–566.

Sanjana, N. E., J. Wright, K. Zheng, O. Shalem, P. Fontanillas, J. Joung, C. Cheng, A. Regev and F. Zhang (2016). “High-resolution interrogation of functional elements in the noncoding genome.” Science 353(6307): 1545–1549.

Sanson, K. R., R. E. Hanna, M. Hegde, K. F. Donovan, C. Strand, M. E. Sullender, E. W. Vaimberg, A. Goodale, D. E. Root, F. Piccioni and J. G. Doench (2018). “Optimized libraries for CRISPR-Cas9 genetic screens with multiple modalities.” Nat Commun 9(1): 5416.

Schmidt, R., Z. Steinhart, M. Layeghi, J. W. Freimer, R. Bueno, V. Q. Nguyen, F. Blaeschke, C. J. Ye and A. Marson (2022). “CRISPR activation and interference screens decode stimulation responses in primary human T cells.” Science 375(6580): eabj4008.

Schmiedel, B. J., C. Gonzalez-Colin, V. Fajardo, J. Rocha, A. Madrigal, C. Ramírez-Suástegui, S. Bhattacharyya, H. Simon, J. A. Greenbaum, B. Peters, G. Seumois, F. Ay, V. Chandra and P. Vijayanand (2022). “Single-cell eQTL analysis of activated T cell subsets reveals activation and cell type-dependent effects of disease-risk variants.” Sci Immunol 7(68): eabm2508.

Schmiedel, B. J., D. Singh, A. Madrigal, A. G. Valdovino-Gonzalez, B. M. White, J. Zapardiel-Gonzalo, B. Ha, G. Altay, J. A. Greenbaum, G. McVicker, G. Seumois, A. Rao, M. Kronenberg, B. Peters and P. Vijayanand (2018). “Impact of Genetic Polymorphisms on Human Immune Cell Gene Expression.” Cell 175(6): 1701–1715.e1716.

Schumann, K., S. S. Raju, M. Lauber, S. Kolb, E. Shifrut, J. T. Cortez, N. Skartsis, V. Q. Nguyen, J. M. Woo, T. L. Roth, R. Yu, M. L. T. Nguyen, D. R. Simeonov, D. N. Nguyen, S. Targ, R. E. Gate, Q. Tang, J. A. Bluestone, M. H. Spitzer, C. J. Ye and A. Marson (2020). “Functional CRISPR dissection of gene networks controlling human regulatory T cell identity.” Nat Immunol 21(11): 1456–1466.

Shifrut, E., J. Carnevale, V. Tobin, T. L. Roth, J. M. Woo, C. T. Bui, P. J. Li, M. E. Diolaiti, A. Ashworth and A. Marson (2018). “Genome-wide CRISPR Screens in Primary Human T Cells Reveal Key Regulators of Immune Function.” Cell 175(7): 1958–1971.e1915.

Simeonov, D. R., B. G. Gowen, M. Boontanrart, T. L. Roth, J. D. Gagnon, M. R. Mumbach, A. T. Satpathy, Y. Lee, N. L. Bray, A. Y. Chan, D. S. Lituiev, M. L. Nguyen, R. E. Gate, M. Subramaniam, Z. Li, J. M. Woo, T. Mitros, G. J. Ray, G. L. Curie, N. Naddaf, J. S. Chu, H. Ma, E. Boyer, F. Van Gool, H. Huang, R. Liu, V. R. Tobin, K. Schumann, M. J. Daly, K. K. Farh, K. M. Ansel, C. J. Ye, W. J. Greenleaf, M. S. Anderson, J. A. Bluestone, H. Y. Chang, J. E. Corn and A. Marson (2017). “Discovery of stimulation-responsive immune enhancers with CRISPR activation.” Nature 549(7670): 111–115.

Skapenko, A., J. Leipe, P. E. Lipsky and H. Schulze-Koops (2005). “The role of the T cell in autoimmune inflammation.” Arthritis Res Ther 7 Suppl 2(Suppl 2): S4–14.

Soskic, B., E. Cano-Gamez, D. J. Smyth, K. Ambridge, Z. Ke, J. C. Matte, L. Bossini-Castillo, J. Kaplanis, L. Ramirez-Navarro, A. Lorenc, N. Nakic, J. Esparza-Gordillo, W. Rowan, D. Wille, D. F. Tough, P. G. Bronson and G. Trynka (2022). “Immune disease risk variants regulate gene expression dynamics during CD4(+) T cell activation.” Nat Genet 54(6): 817–826.

Soskic, B., E. Cano-Gamez, D. J. Smyth, W. C. Rowan, N. Nakic, J. Esparza-Gordillo, L. Bossini-Castillo, D. F. Tough, C. G. C. Larminie, P. G. Bronson, D. Willé and G. Trynka (2019). “Chromatin activity at GWAS loci identifies T cell states driving complex immune diseases.” Nat Genet 51(10): 1486–1493.

Squair, J. W., M. Gautier, C. Kathe, M. A. Anderson, N. D. James, T. H., Hutson, R., Hudelle, T. Qaiser, K. J. E. Matson, Q. Barraud, A. J. Levine, G. La Manno, M. A. Skinnider and G. Courtine (2021). “Confronting false discoveries in single-cell differential expression.” Nat Commun 12(1): 5692.

Trynka, G. and S. Raychaudhuri (2013). “Using chromatin marks to interpret and localize genetic associations to complex human traits and diseases.” Curr Opin Genet Dev 23(6): 635–641.

Ulirsch, J. C., C. A. Lareau, E. L. Bao, L. S. Ludwig, M. H. Guo, C. Benner, A. T. Satpathy, V. K. Kartha, R. M. Salem, J. N. Hirschhorn, H. K. Finucane, M. J. Aryee, J. D. Buenrostro and V. G. Sankaran (2019). “Interrogation of human hematopoiesis at single-cell and single-variant resolution.” Nat Genet 51(4): 683–693.

Umans, B. D., A. Battle and Y. Gilad (2021). “Where Are the Disease-Associated eQTLs?” Trends Genet 37(2): 109–124.

van der Wijst, M. G. P., H. Brugge, D. H. de Vries, P. Deelen, M. A. Swertz and L. Franke (2018). “Single-cell RNA sequencing identifies celltype-specific cis-eQTLs and co-expression QTLs.” Nat Genet 50(4): 493–497.

Vujkovic, M., J. M. Keaton, J. A. Lynch, D. R. Miller, J. Zhou, C. Tcheandjieu, J. E. Huffman, T. L. Assimes, K. Lorenz, X. Zhu, A. T. Hilliard, R. L. Judy, J. Huang, K. M. Lee, D. Klarin, S. Pyarajan, J. Danesh, O. Melander, A. Rasheed, N. H. Mallick, S. Hameed, I. H. Qureshi, M. N. Afzal, U. Malik, A. Jalal, S. Abbas, X. Sheng, L. Gao, K. H. Kaestner, K. Susztak, Y. V. Sun, S. L. DuVall, K. Cho, J. S. Lee, J. M. Gaziano, L. S. Phillips, J. B. Meigs, P. D. Reaven, P. W. Wilson, T. L. Edwards, D. J. Rader, S. M. Damrauer, C. J. O’Donnell, P. S. Tsao, K. M. Chang, B. F. Voight and D. Saleheen (2020). “Discovery of 318 new risk loci for type 2 diabetes and related vascular outcomes among 1.4 million participants in a multi-ancestry meta-analysis.” Nat Genet 52(7): 680–691.

Wakefield, J. (2009). “Bayes factors for genome-wide association studies: comparison with P-values.” Genet Epidemiol 33(1): 79–86.

Wallace, C. G., C.; Plagnol V. (2022). “coloc: Colocalisation Tests of Two Genetic Traits.” from https://cran.r-project.org/web/packages/coloc/index.html.

Xie, S., J. Duan, B. Li, P. Zhou and G. C. Hon (2017). “Multiplexed Engineering and Analysis of Combinatorial Enhancer Activity in Single Cells.” Molecular cell 66(2): 285–299.e285.

Yang, J., T. Ferreira, A. P. Morris, S. E. Medland, P. A. Madden, A. C. Heath, N. G. Martin, G. W. Montgomery, M. N. Weedon, R. J. Loos, T. M. Frayling, M. I. McCarthy, J. N. Hirschhorn, M. E. Goddard and P. M. Visscher (2012). “Conditional and joint multiple-SNP analysis of GWAS summary statistics identifies additional variants influencing complex traits.” Nat Genet 44(4): 369–375, s361-363.

Yang, J., S. H. Lee, M. E. Goddard and P. M. Visscher (2011). “GCTA: a tool for genome-wide complex trait analysis.” Am J Hum Genet 88(1): 76–82.

Yao, D. W., L. J. O’Connor, A. L. Price and A. Gusev (2020). “Quantifying genetic effects on disease mediated by assayed gene expression levels.” Nat Genet 52(6): 626–633.

Yazar, S., J. Alquicira-Hernandez, K. Wing, A. Senabouth, M. G. Gordon, S. Andersen, Q. Lu, A. Rowson, T. R. P. Taylor, L. Clarke, K. Maccora, C. Chen, A. L. Cook, C. J. Ye, K. A. Fairfax, A. W. Hewitt and J. E. Powell (2022). “Single-cell eQTL mapping identifies cell type-specific genetic control of autoimmune disease.” Science 376(6589): eabf3041.

Young, A. M. H., N. Kumasaka, F. Calvert, T. R. Hammond, A. Knights, N. Panousis, J. S. Park, J. Schwartzentruber, J. Liu, K. Kundu, M. Segel, N. A. Murphy, C. E. McMurran, H. Bulstrode, J. Correia, K. P. Budohoski, A. Joannides, M. R. Guilfoyle, R. Trivedi, R. Kirollos, R. Morris, M. R. Garnett, I. Timofeev, I. Jalloh, K. Holland, R. Mannion, R. Mair, C. Watts, S. J. Price, P. J. Kirkpatrick, T. Santarius, E. Mountjoy, M. Ghoussaini, N. Soranzo, O. A. Bayraktar, B. Stevens, P. J. Hutchinson, R. J. M. Franklin and D. J. Gaffney (2021). “A map of transcriptional heterogeneity and regulatory variation in human microglia.” Nat Genet 53(6): 861–868.

Zhernakova, D. V., P. Deelen, M. Vermaat, M. van Iterson, M. van Galen, W. Arindrarto, P. van ‘t Hof, H. Mei, F. van Dijk, H. J. Westra, M. J. Bonder, J. van Rooij, M. Verkerk, P. M. Jhamai, M. Moed, S. M. Kielbasa, J. Bot, I. Nooren, R. Pool, J. van Dongen, J. J. Hottenga, C. D. Stehouwer, C. J. van der Kallen, C. G. Schalkwijk, A. Zhernakova, Y. Li, E. F. Tigchelaar, N. de Klein, M. Beekman, J. Deelen, D. van Heemst, L. H. van den Berg, A. Hofman, A. G. Uitterlinden, M. M. van Greevenbroek, J. H. Veldink, D. I. Boomsma, C. M. van Duijn, C. Wijmenga, P. E. Slagboom, M. A. Swertz, A. Isaacs, J. B. van Meurs, R. Jansen, B. T. Heijmans, P. A. t Hoen and L. Franke (2017). “Identification of context-dependent expression quantitative trait loci in whole blood.” Nat Genet 49(1): 139–145.

Zhu, J., H. Yamane and W. E. Paul (2010). “Differentiation of effector CD4 T cell populations (*).” Annu Rev Immunol 28: 445–489.

Zimmerman, K. D., M. A. Espeland and C. D. Langefeld (2021). “A practical solution to pseudoreplication bias in single-cell studies.” Nat Commun 12(1): 738.

